# A Modular Genetic Toolbox for Precise Gene Regulation and Multi-Color Imaging in Streptococci

**DOI:** 10.1101/2025.05.10.653270

**Authors:** Johann Mignolet, Johan Staub, Jan Roelof van der Meer, Jan-Willem Veening, Virginie Libante

## Abstract

Fluorescent labeling is a powerful tool in microbiology allowing live cell imaging and providing insights into dynamic cellular processes, quantification of gene expression and protein subcellular localization. Although multicolor imaging is widely used in *Streptococcus pneumoniae* and *S. mutans*, variants other than the green fluorescent protein (GFP) have rarely been applied in other streptococcal species. To address this gap in the streptococcal molecular toolbox, we benchmarked five different fluorescent proteins. The various fluorescent proteins were fused to the C-terminus of *S. pneumoniae* HlpA, a small non-specific DNA binding histone-like protein. These reporters, combined with four different antibiotic resistance genes, were engineered with various expression systems (inducible or constitutive) to form versatile cassettes. We provide methods to transfer these cassettes to different streptococcal species including *S. salivarius* and *S. thermophilus*. As a proof of concept, we generated a triple labeled *S. salivarius* strain in which HlpA, FtsZ and DivIVA were fused to three spectrally-distinct compatible fluorescent proteins. Multiple fluorescent labeling has broad applications for deciphering a wide range of scientific problems, from cellular processes to infectious disease mechanisms. The availability of these cassettes should allow for a wider use of single-cell labeling strategies in the Streptococcus clade and other closely related bacteria.

## INTRODUCTION

Since the mid-1990s, the use of the green fluorescent protein (GFP) has been a powerful biomolecular tool for gene expression, protein localization/tracking and live cell detection [1, 2]. In bacteria, starting with *Escherichia coli*, *Bacillus subtilis* and *Pseudomonas putida*, GFP has been successfully used to study protein and DNA localization [3, 4]. GFP variants have been selected to solve problems of low intensity, folding, solubility and inclusion bodies at elevated temperatures [5]. In parallel, mutant proteins with different excitation and emission spectra and fluorescence intensity were discovered or engineered to allow for multiple dynamic labeling of bacterial communities [6], or proteins [7].

The human pathogen *Streptococcus pneumoniae* is a Gram-positive bacterial species for which many molecular tools have been developed. Extensive work has been done on *S. pneumoniae* to develop labeling strategies, and a collection of compatible fluorescent proteins (FPs) for multiple and dynamic labeling optimized for super-resolution microscopy have been made available (listed in [8, 9]). The benefits of multi-labeling could be two-fold. It allows to (i) discriminate between several populations of cells/species labeled with compatible fluorescent proteins (for instance with *Streptococcus mutans* [10, 11]), or (ii) simultaneously track the subcellular location of numerous proteins in a single cell, as shown with integrative vectors [9]. Despite these studies, the implementation of labeling strategies in other streptococcal species has remained limited [10–16].

Sufficient and adequate abundance of FPs is a prerequisite for efficient population or cell monitoring. To stabilize fluorescent proteins, translational fusions to highly abundant and stable proteins have been engineered [1]. In *S. pneumoniae*, the histone-like protein HlpA (a.k.a. HU) fused to an FP at its C-terminal end, gave remarkably brightly labeled cells [17]. This was due to (i) the stable chromosomal integration of the fusion gene downstream of native *hlpA*, (ii) the high transcript levels of *hlpA* throughout all stages of the cell cycle, and (iii) the nucleoid association of the fluorescent signal, without deleterious effects, which limits the diffusion of the FP. This approach was first applied to *S. pneumoniae* for *in vivo* live cell imaging of two different HlpA-FP labeled strains within mouse infection experiments [17] and then, for biofilm formation with two combined strains [18]. HlpA is essential for cell survival in streptococci [19–21], and is largely conserved across prokaryotes [22, 23], allowing this labeling strategy to be applied to a wider range of bacteria.

In parallel to protein stability as such, the system and timing of expression is of major importance. Constitutive production of (recombinant) proteins can have drastic effects on cell physiology, slowing down growth rate or provoking abnormal cell morphologies [24, 25]. The overload of overproduced protein can consume the energy and nutrients of a cell and influence its metabolic state. Moreover, inappropriate timing of expression can collide with the cell cycle. Inducible expression systems offer a solution to minimize collateral toxicity from protein overproduction. Most studied promoters from lactic acid bacteria are leaky, trigger too strong expression or are not sufficiently characterized [26–29]. In beneficial streptococci, some inducible promoters have been adapted for expression studies (*e.g.* P*lac*, P*xyl*, and P*tet*) [30–33]. Nonetheless, such controllable tools are lacking in pathogenic streptococci, apart from *S. pneumoniae and Streptococcus pyogenes* [16] and lactic acid bacteria in general.

The aim of this work is to expand and implement the molecular toolbox for streptococcal species and, to a broader scope, for lactic acid bacteria. Here, we make available FP-tagged HlpA^Sp^ constructs associated with four antibiotic resistances usable for streptococci (*hlpA^Sp^-fp*- Ab^R^ cassettes). The selected FPs (mTurquoise2, mNeonGreen, a variant of msfYFP, mScarlet- I and mKate2) allow simultaneous tripartite detection. To have the broadest application, the FP-fusion cassettes were cloned in *Escherichia coli* on a high-copy number plasmid (pJet1.2/blunt, Thermo Scientific [34]) with a non-functional promoter sequence, because expression of *Streptococcus intermedius hlpA* is toxic in *E. coli* [19]. The cassettes can be used for overlapping PCR engineering prior to chromosomal integration into naturally competent streptococcal strains or on shuttle vectors. As a proof of principle, FPs were fused to proteins exhibiting specific subcellular localization (in addition to HlpA) and co-produced from endogenous, constitutive or inducible promoters. The molecular toolbox presented here will facilitate single cell analysis and gene expression studies in streptococci.

## RESULTS

### Chromosomal insertion at the hlpA locus of Streptococcus thermophilus

We selected the following five fluorescent proteins (FPs) for expression in streptococcus: mTurquoise2, mNeonGreen, msfYFP, mScarlet-I and mKate2. Genes encoding the FPs were previously codon-optimized for *S. pneumoniae* and shown to be well expressed in *S. pneumoniae* when fused at the C-terminus of HlpA [8, 17]. The five *hlpA^Sp^-fp* fusions were amplified as PCR products and inserted into the pJet1.2/blunt positive selection vector. After electrotransformation in *E. coli*, we validated by sequencing that all vectors carried the correct *hlpA^Sp^-fp* insert downstream of the T7 RNA polymerase promoter. For selection and maintenance, resistance genes for chloramphenicol (*cat*Q; Cm^R^), erythromycin (*erm*(B); Ery^R^), kanamycin (APH(3’)-IIIa; Kan^R^), and spectinomycin (ANT(9); Spec^R^) were cloned with their own promoters (functional in both *E. coli* and firmicutes) downstream of the *hlpA^Sp^-fp* fragments, ensuring the same transcriptional orientation (Figure 1a). The constructed plasmids (available at Addgene #206810-206829) are replicative only in *E. coli* (Supplementary Table S1) and can be used as templates for PCR amplification of *hlpA^Sp^-fp* or *hlpA^Sp^-fp*-Ab^R^ cassettes (primer sequences listed in the Supplementary Table S2).

**Figure 1.**
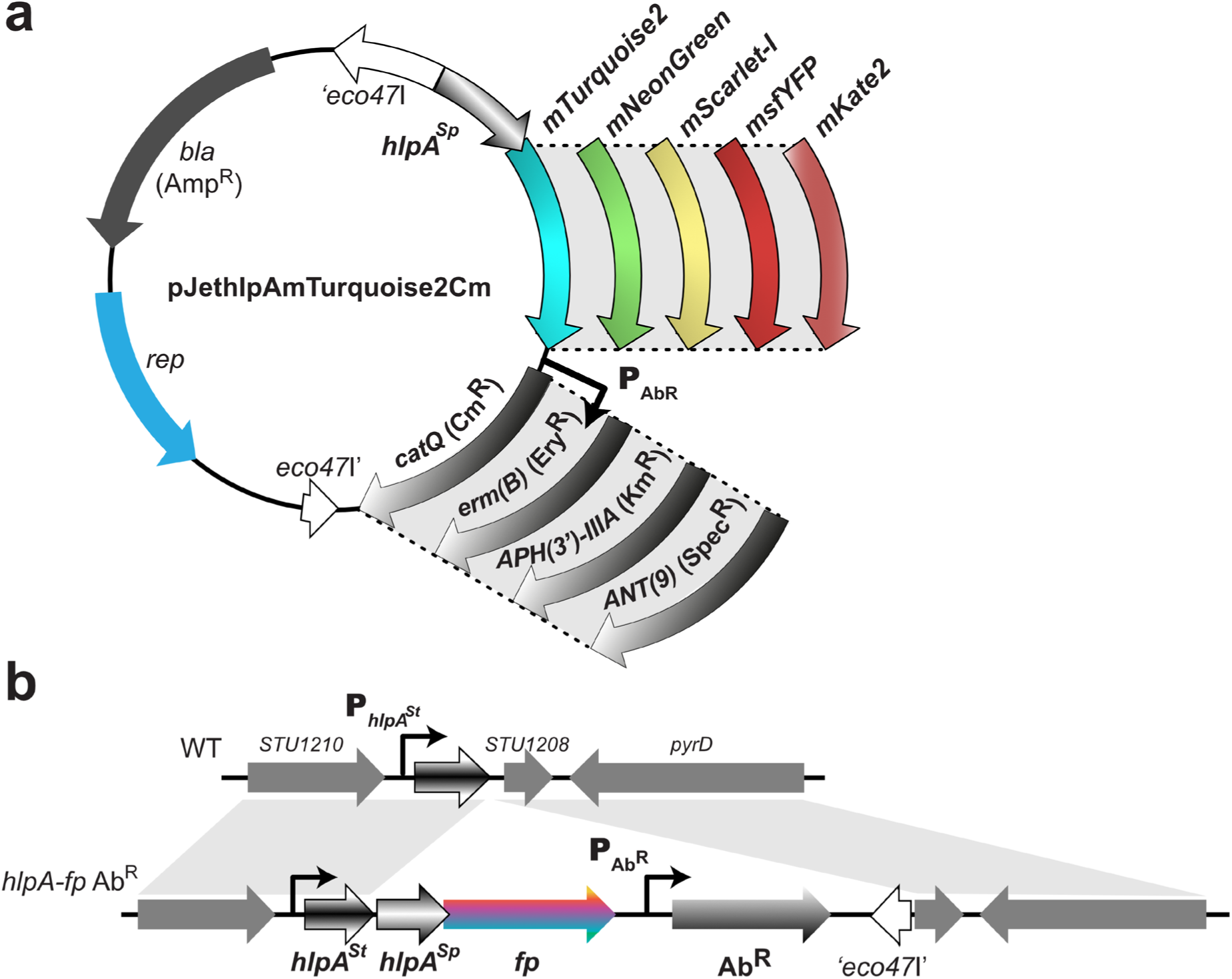
Engineering toolbox for gene insertion and fluorescent protein production in *S. thermophilus* chromosome. (a) General scheme of the pJethlpAmTurquoise2Cm and its derivatives. The plasmid includes the pBR322 origin (in blue) for propagation in *E. coli*. The ampicillin resistance gene (*bla*) is shown in grey. The *eco47*I gene (in white) is disrupted by the *hlpA^Sp^-fp* fusion, the antibiotic resistance genes (Ab^R^) cloned under their own promoter control and transcriptional terminators. A set of 20 plasmids was created, each containing a different *hlpA^Sp^* chimeric gene linked to one out of five genes encoding fluorescent protein (mTurquoise2, mNeonGreen, msfYFP, mScarlet-I and mKate2) and one out of four antibiotic resistance genes (*catQ*, *erm*(B), APH(3’)-IIIa, ANT(9)). (b) Homologous recombination at the *hlpA* locus of *S. thermophilus*. On the top panel, the recipient chromosomal locus of *S. thermophilus* LMG18311, *hlpA^St^* for *hlpA^Sp^*-*fp-* Ab^R^ cassette integration is shown. On the bottom panel, the final insertion product is depicted. After homologous recombination, the *hlpA^Sp^*-*fp-*Ab^R^ cassette is inserted downstream of the *hlpA^St^* gene, as highlighted by the homologous gray regions. Promoters are marked with arrows symbols.

The *S. thermophilus hlpA* chromosomal locus was previously characterized by Dixon-Fyle and Caro [35] (Figure 1b). Insertion of the *hlpA^Sp^-fp-*Ab^R^ was performed following the strategy outlined by Kjos et al [17]. Since *hlpA* is essential in *S. pneumoniae* D39, *hlpA^Sp^-fp* was integrated downstream of the native *hlpA^St^* gene. Natural competence transformation combined with antibiotic selection using the *hlpA^Sp^-fp-*Ab^R^ cassettes, provided an efficient and time-saving method for strain modification (Figure 1b, successful recombination events shown in gray). Transcription of *hlpA^Sp^-fp* was driven from the native *hlpA^St^* promoter, forming a single transcriptional unit with a duplication of the ribosomal binding site of *hlpA^St^*. Clones were verified by PCR and sequencing.

### Quantitative analysis of fluorescence detection in growing cultures of S. thermophilus hlpA^Sp^-fp

To assess the fluorescence profiles of each FP fusion, cultures in early stationary phase were diluted in fresh M17L medium to a starting culture turbidity (OD at 590 nm) of 0.02. Growth and fluorescence signals were recorded every 10 minutes during 24 hours in a microtiter plate assay (see Methods). Growth curves of the engineered strains were comparable to the parental LMG18311 strain, indicating that the integration of *hlpA^Sp^-fp* downstream of the *hlpA^St^* gene had no or minimal effect on growth (Figure 2 and Supplementary Figure S1). No detectable fluorescence could be observed for mTurquoise2 and mKate2 in this medium and with our available microtiter plate setup (Supplementary Figure S1). However, high signals in the channels for mNeonGreen, msfYFP, and mScarlet-I fluorescence were detected during growth (Figure 2a-c). Fluorescence levels continuously increased with cell density until the bacterial population reached the stationary phase. The signal peaked early in stationary phase for HlpA^Sp^-mNeonGreen (Figure 2a) and HlpA^Sp^-msfYFP (Figure 2b), and approximately 2 hours later for HlpA^Sp^-mScarlet-I, suggesting a longer maturation time of mScarlet-I in *S. thermophilus* (Figure 2c), as previously reported for eucaryote and other bacterial expression systems [36]. Approximately 10 hours after this peak value of fluorescence, the levels declined to the background level of the wild-type LMG18311 culture. There was some overlap between the HlpA^Sp^-msfYFP and HlpA^Sp^-mNeonGreen fusion protein fluorescence emissions (Supplementary Figure S2). However, fluorescence measurements in relative fluorescent units (RFUs) for specific excitation and emission wavelengths of each protein showed a stronger signal in their selective channel. No cross-detection was observed between HlpA^Sp^-mScarlet-I and HlpA^Sp^-mNeonGreen or HlpA^Sp^-msfYFP suggesting that mScarlet-I and mNeonGreen/msfYFP can be simultaneously recorded in microtiter plate assays (Supplementary Figure S2). Since the integration locus, promoter, and protein fusion design were identical for the three fluorescent proteins, the observed fluorescence curves likely reflect differences in the folding kinetics and stability of the HlpA^Sp^-FP fusions, which are directly influenced by the specific fluorescent proteins.

**Figure 2.**
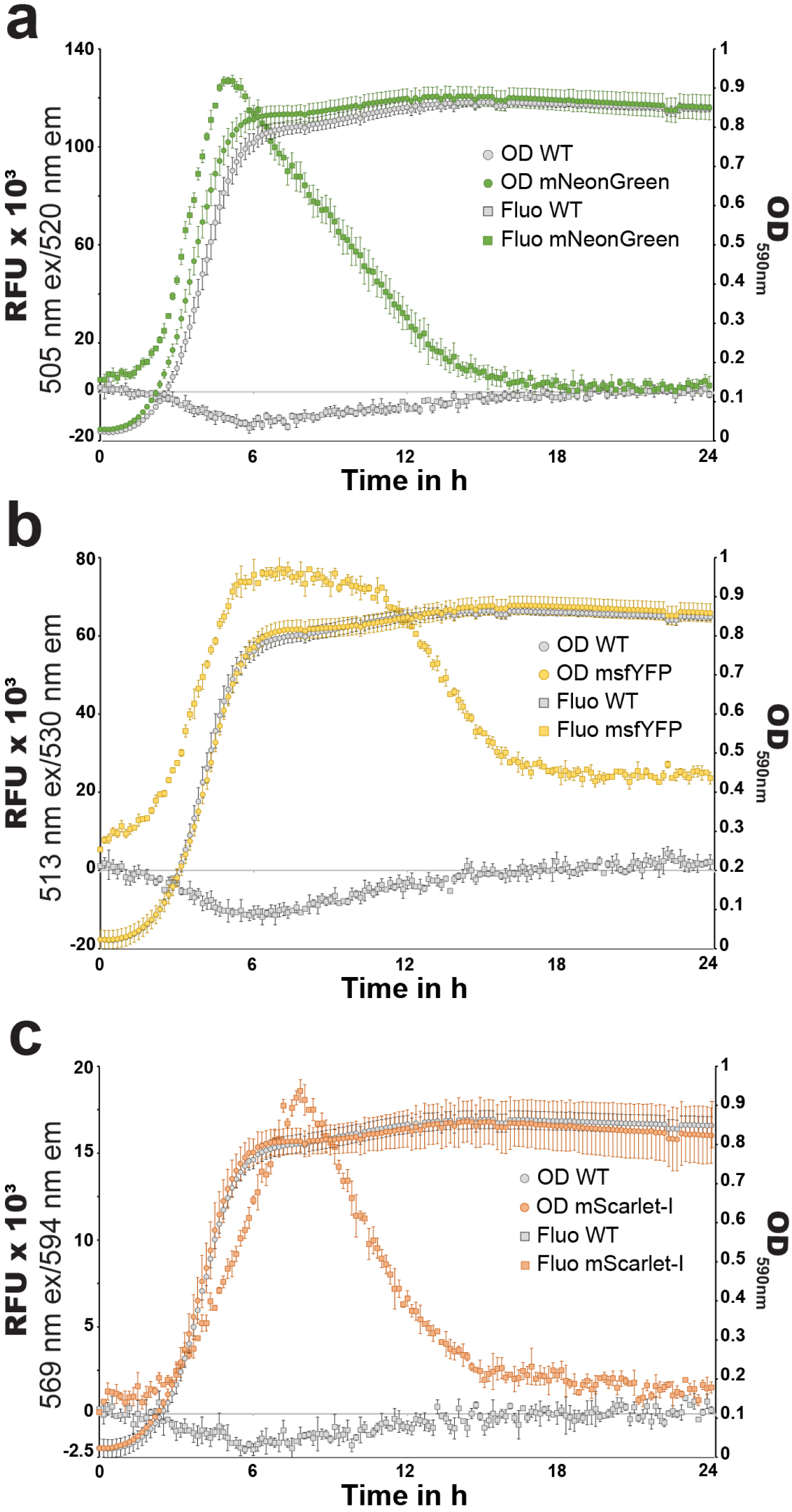
Fluorescence quantification of *S. thermophilus hlpA^Sp^-fp* during growth. Fluorescence measurements of growing *S. thermophilus* LMG18311 (WT) and its derivative strains in M17L medium. Growth curves are assessed via culture turbidity at 590 nm (round symbols) and fluorescence measurements are expressed in Relative Fluorescence Units (RFUs, after subtracted the M17L signal, square symbols). LMG18311 derivatives shown by colored symbols, while background fluorescence in the parental LMG18311 strain is plotted in grey. RFU measurements were taken at 505 nm ex/520 nm em (mNeonGreen), 513 nm ex/530 nm em (msfYFP), and 569 nm ex/594 nm em (mScarlet-I). Due to significant background fluorescence for the M17L medium, the parental strain LMG18311 in m17L reduced the signal, resulting in negative values when the blank (M17L without cells) was subtracted (grey squares). Data points are the means from triplicate assays with error bars indicating the mean ± standard deviation.

### Microscopy-based quantification of fluorescent signal in cultures of S. thermophilus producing HlpA^Sp^-FP

Epifluorescence microscopy of *S. thermophilus* cells producing various HlpA^Sp^-FP fusions allowed us to define the signal-to-noise ratio (mean intensity of the cellular area divided by the mean intensity of the non-cell background area in a specific channel) for each FP with its respective filter. Light source intensity and exposure time were adjusted to minimize photobleaching for each individual fusion. As shown in Figure 3a and 3b, the mTurquoise2, mNeonGreen, msfYFP and mScarlet-I HlpA fusions produced a high fluorescence signal that was restricted to the bacterial nucleoid, as expected for a non-specific nucleoid binding protein, while the signal associated with the mKate2 fusion was dimmer. Moreover, recording the fluorescence values and evaluating the signal-to-noise ratio of each fusion for each set of filters showed that the HlpA^Sp^-mTurquoise2 fusion is fully compatible with every other fusion (no overlap between channels) in *S. thermophilus* (Figure 3b). By contrast, two pairs of FPs, HlpA^Sp-^mNeonGreen/HlpA^Sp^-msfYFP and HlpA^Sp^-mScarlet-I/HlpA^Sp^-mKate2, could not be differentiated specifically and were incompatible in our microscope setup. However, their signals do not bleed through other channels, meaning they can be imaged with all other FPs tested here. To conclude, the nucleoid-associated signal of HlpA^Sp^-FP fusions (rather than a diffuse cytoplasmic fluorescence for FPs alone) enables clear identification of cells even at low signal intensities. In *S. thermophilus*, HlpA^Sp^-mTurquoise2, HlpA^Sp^-msfYFP and HlpA^Sp^-mScarlet-I emit distinct and compatible fluorescent signals, when detected with their respective filters, allowing accurate identification and enumeration of individual cells in a mixed population (Figure 4).

**Figure 3.**
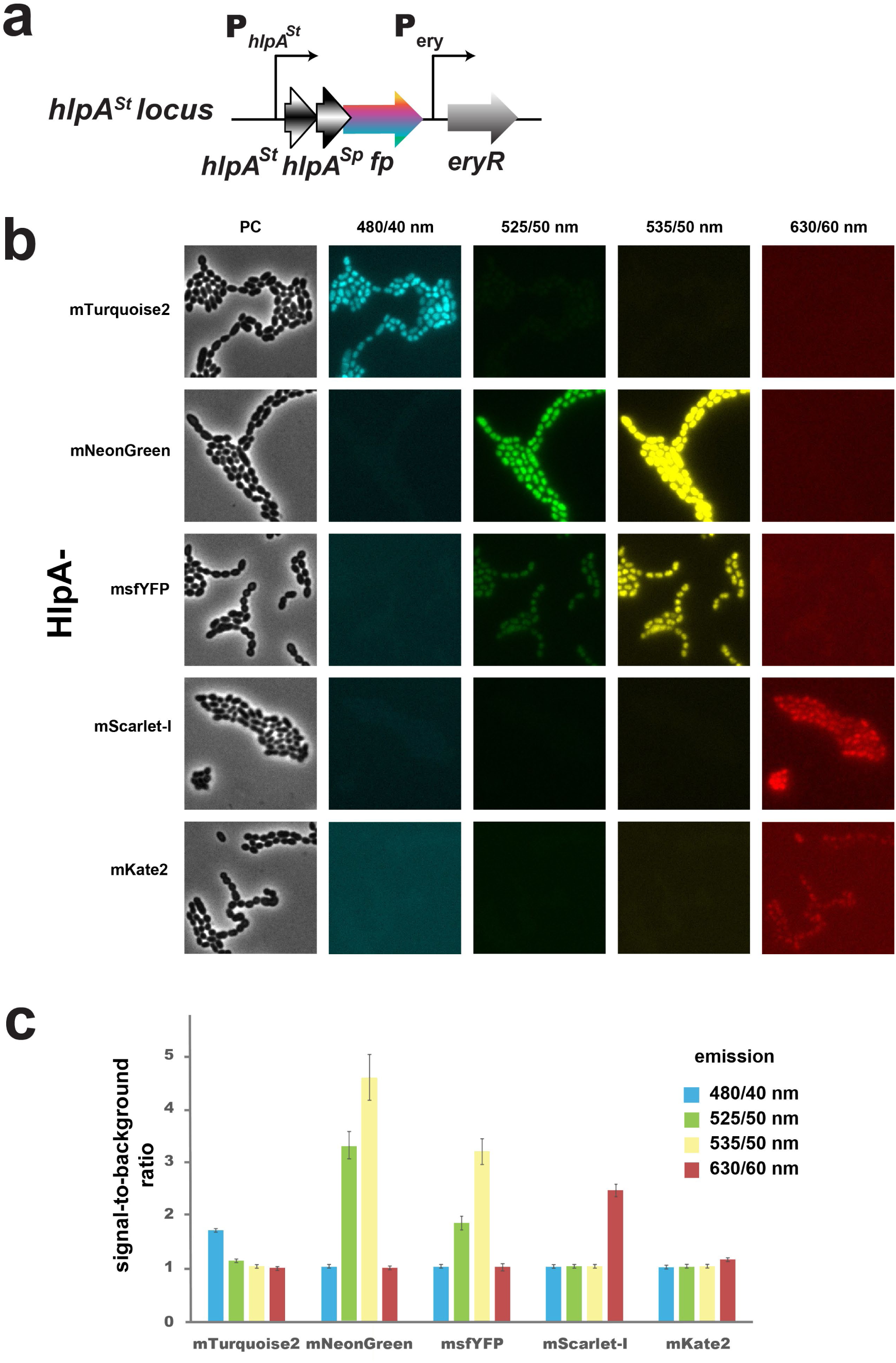
Single-cell visualization and quantification of FP signals in *S. thermophilus*. (a) Genetic structure of the *hlpA-fp* integrated at the native *hlpA^St^* locus under the control of its own promoter. The Ery^R^ gene is used to select chromosomal integration of *hlpA-fp* constructs. (b) Cell micrographs of *S. thermophilus hlpA^Sp^-fp* expressing strains for HlpA^Sp^-mTurquoise2, HlpA^Sp^-mNeonGreen, HlpA^Sp^-msfYFP, HlpA^Sp^-mScarlet-I or HlpA^Sp^-mKate2 (as indicated on the left) in the respective fluorescence channels, or in phase contrast (PC). Images are scaled to same brightness and contrast. Scale bars represent 2 µm. (c) Signal-to-noise ratio for the five FPs of panel (b) in *S. thermophilus* in their cognate emission channels: 480/40 nm (blue), 525/50 nm (green), 535/50 nm (yellow), and 630/60 nm (red). Cells were segmented with Fiji to quantify the mean fluorescence pixel intensity per cell, which is divided by the mean pixel intensity of image background (excluding cells). Bars represent the means ± standard error from three images.

**Figure 4.**
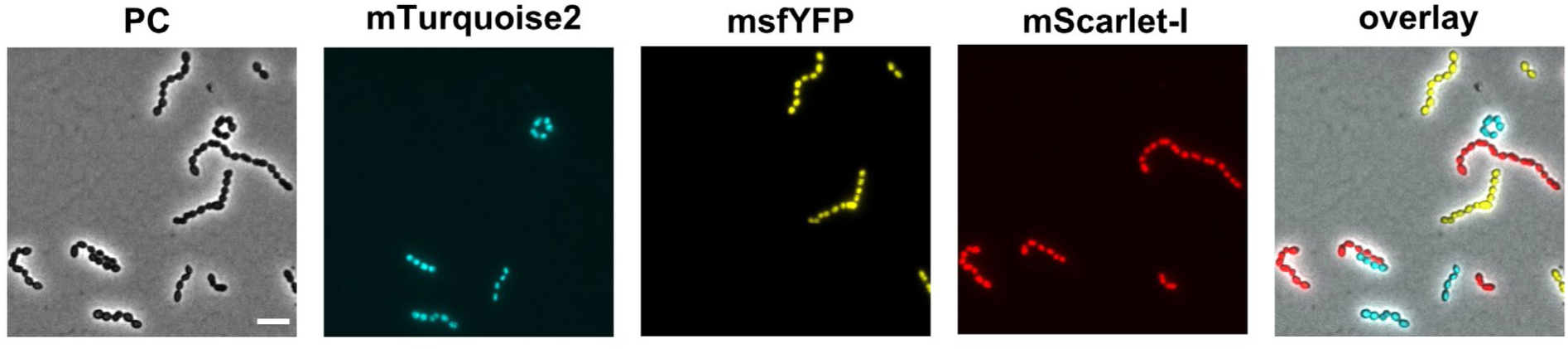
Fluorescence-based discrimination of mixed *S. thermophilus* populations producing HlpA-FP Fusions. Cell micrographs show an equal proportions mixture of three *S. thermophilus* LMG18311 populations expressing HlpA^Sp^-mTurquoise2, HlpA^Sp^-msfYFP or HlpA^Sp^-mScarlet-I, imaged in phase contrast (PC), or by the specific fluorescence settings for each of the FPs (485/40 nm/blue, 535/50 nm /yellow, and 630/60 nm/red), along with a color overlay image. Scale bars represent 2 µm.

### Constitutive hlpA^sp^-fp expression from ectopic locus in Streptococcus salivarius

To validate that the HlpA^Sp^-FP fusions are exploitable in other species, we adapted the system to *Streptococcus salivarius*, a human commensal that is highly prevalent in the digestive tract. *S. salivarius* is genetically amenable due to its natural competence but fluorescent tools to perform single-cell analyses and monitor population dynamics are missing. To probe the versatility of our genetic toolbox, we inserted the *hlpA^Sp^-fp* constructs at a non-essential ectopic locus in *S. salivarius* next to *tRNA^Ser^*and expressed them under the control of the constitutive P32 promoter [32] (Supplementary Figure S3a). The mTurquoise2, mNeonGreen, msfYFP, mScarlet-I, and mKate2 fusions produced a strong homogeneous signal localized to the nucleoid (Figure 5a and Supplementary Figure S3b), well distinguishable above background (Supplementary Figure S3c). Moreover, most pairs are compatible for multi-labeling, and the trio mTurquoise2/mNeonGreen/mScarlet-I enables simultaneous imaging with high signal-to-noise ratio and minimal crosstalk in *S. salivarius* on a standard widefield epifluorescence microscope.

**Figure 5.**
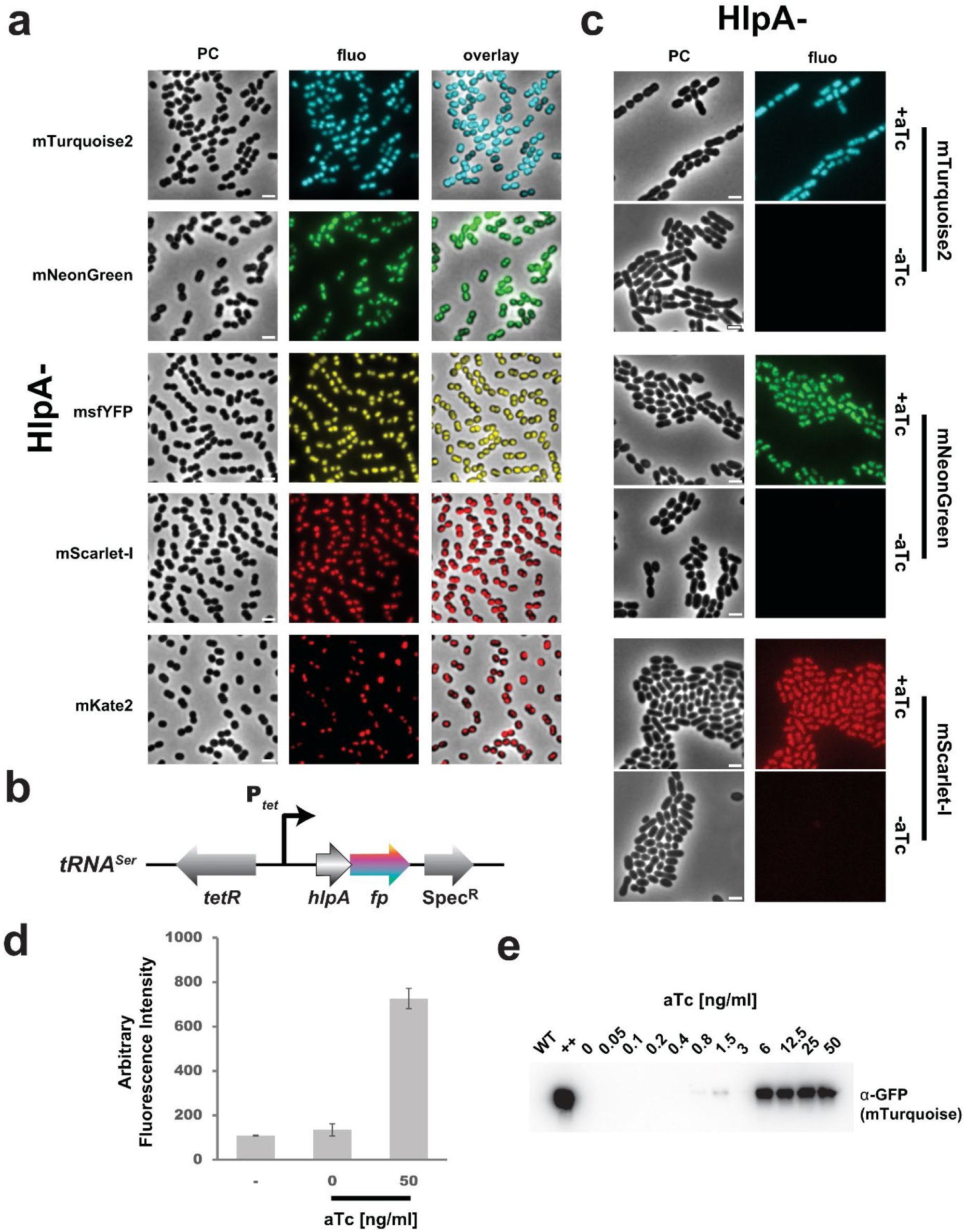
Constitutive and inducible nucleoid-FP labeling in *S. salivarius*. (a) Micrographs of *S. salivarius* strains expressing the HlpA^Sp^ nucleoid-associated factor FP fusion for each of the fluorescent protein variants, as indicated on the left, imaged by phase contrast (PC), or fluorescence recorded in the cognate channel, and an overlay. The scale bar represents 2 μm. (b) Scheme of the genetic structure of the P*tet*-inducible *hlpA-fp* integrated at the *tRNA^Ser^* locus. The *tetR* repressor gene, with its own promoter (not depicted here), points in the opposite orientation compared to the P*tet*. The Spec^R^ gene is used to select chromosomal integration of *hlpA-fp* constructs. (c) Single-cell imaging of aTc-inducible tagged HlpA-FP with mTurquoise2, mNeonGreen or mScarlet-I in *S. salivarius* cells, incubated with (+) or without (-) 50 ng.ml^-1^ aTc for 2 hours. All images are scaled to the same brightness. The scale bar equals 2 μm. (d) Mean single cell fluorescence of the aTc-induced (50 ng.ml^-1^) or uninduced (0) *hlpA^Sp^-mTurquoise2* strain of *S. salivarius*, in comparison to the wild-type non-labeled HSISS4 strain as a negative control (-). Bars show the mean ± standard error from cells segmented on three separate images. (e) Anti-GFP antibody stained immunoblot showing the HlpA-mTurquoise2 levels upon growth in M17G as a function of aTc concentration (from 0.05 to 50 ng.ml^-1^). WT, a non-labeled control strain; (++), a constitutive overexpression mutant (expression from P32).

### Ectopic chromosomal expression under inducible promoter control in *Streptococcus salivarius*

Misexpression at an inappropriate time of the cell cycle or cell growth might turn out to have toxic side effects. Therefore, we evaluated several inducible promoters for FP expression in *S. salivarius* that could be operated simultaneously. Two xylose-responsive promoters with distinct strength and leakiness have previously been reported for *S. salivarius* [32]. As further inducible promoter we adapted the tetracycline-responsive promoter PT5-3 (hereafter referred to as P*tet*) that was optimized for tightness and strength in *S. pneumoniae* [37]. Transformation products were assembled by overlapping PCRs to replace the P32 promoter with P*tet* upstream of the mTurquoise2, mNeonGreen or mScarlet-I fusion genes at the *tRNA^Ser^* locus (Figure 5b). Induction for 4h30 with a saturating concentration of anhydrotetracycline (aTc; 50 ng.ml^-1^) to derepress P*tet* caused a strong, homogeneous signal under the microscope for each of the three constructs (Figure 5c). In contrast, cells grown in the absence of aTc emitted fluorescence close to the non-cell background. Quantification of P*tet*-*hlpA^Sp^-*mNeonGreen fluorescence under aTc-induction showed 5.4-fold increase compared to uninduced cells (Figure 5d). Western blotting confirmed induction of HlpA^Sp^-mTurquoise2 in *S. salivarius* as a function of aTc concentrations (Figure 5e), saturating above an aTc concentration of 6 ng ml^-1^ and decreasing to background in a non-mTurquoise2 WT strain at 0.4 ng. ml^-1^. This range of aTc concentrations is similar to induction curves previously reported for *S. pneumoniae* [37]. However, in comparison to the same protein fusion under constitutive P32 expression (++; Figure 5e), the level of HlpA^Sp^-mTurquoise2 from P*tet* at saturating concentrations of aTc was markedly lower. In conclusion, the aTc-inducible promoter in *S. salivarius* allows a titratable, robust, and mild-to-strong protein expression. Initially implemented in *S. pneumoniae*, it can be transferred to other phylogenetically distant streptococci such as *S. salivarius*, and possibly outside of the *Streptococcus* genus.

### Fluorescently-tagged proteins to image and track subcellular localization in S. salivarius

We wondered if we could exploit mTurquoise2, mNeonGreen, and/or mScarlet-I fusions to follow protein dynamics at the single-cell level in *S. salivarius*. A highly dynamic and nearly universal landmark for the future site of division in bacteria is the so-called Z-ring formed by the tubulin-like protein FtsZ [38]. To image the division site in *S. salivarius*, we engineered strains to produce FP-tagged fusions of *S. salivarius* FtsZ. Considering that constitutive overexpression of *ftsZ* is likely to alter the Z-ring polymerization state and might be deleterious to cell shape [39], we used the *tRNA^Ser^* platform to ectopically integrate *ftsZ-fp* under the inducible P*tet* control as a second copy of *ftsZ* (Figure 6a). As shown in Figure 6b, all FtsZ-FP fusions formed well defined rings at the midcell, in line with what was observed for *S. pneumoniae* [40–42]. As cells entered the late phase of cytokinesis, the FtsZ fluorescent signal faded away from the equatorial plane and new Z-rings appeared at the quarter positions representing the future division sites (Movie S1). These experiments validate that *S. salivarius* has a division mode similar to *S. pneumoniae* in which cells elongate and divide in a single plane, in contrast to true cocci like *Staphylococcus aureus* [43].

**Figure 6.**
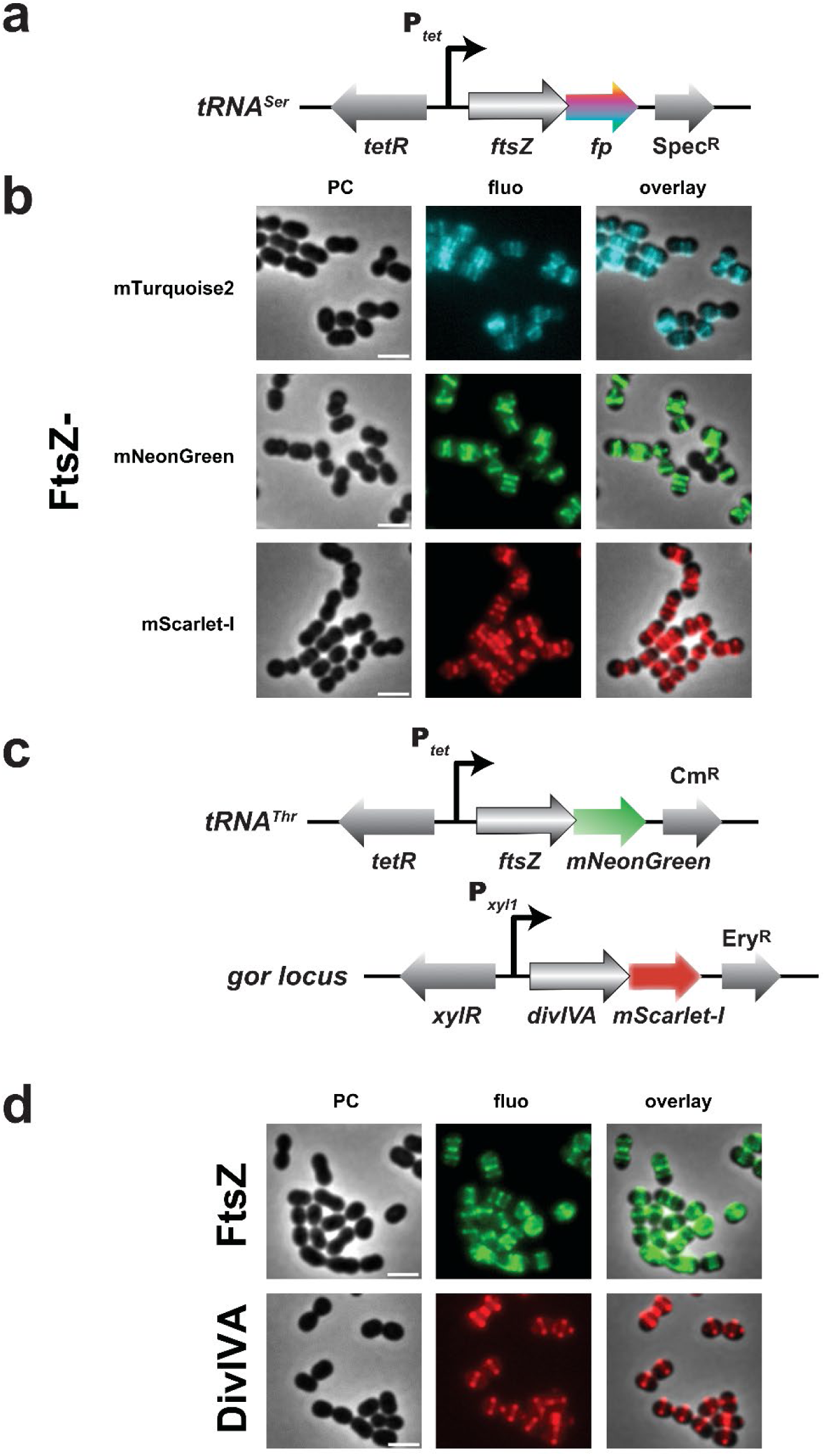
Subcellular localization of FtsZ and DivIVA in *S. salivarius*. (a) Genetic structure of the P*tet*-inducible *ftsZ-fp* integrated at the *tRNA^Ser^* locus. (b) Micrographs of single cells of aTc-induced FtsZ-FP (mTurquoise2, mNeonGreen or mScarlet-I; 50 ng ml^-1^ aTc for 2 hours before imaging). Label indications and scale bar as before (2 μm). (c) Genetic structure of the P*tet*-inducible *ftsZ-mNeonGreen* integrated at the *tRNA^Thr^* locus and the P*xyl1*-inducible *divIVA-mScarlet-I* integrated at the *gor* locus in *S. salivarius*. (d) Micrographs of single cell aTc-derepressed FtsZ-mNeonGreen and xylose-induced DivIVA-mScarlet-I fusions, after incubation with the inducers for 2 hours before imaging. The scale bar equals 2 μm.

### Monitoring protein choreography in real-time with multi-labeling

To monitor the interplay between two or more proteins, multicolor labeling can be used to assess protein localization simultaneously in the same cell. We thus wanted to create a strain with multiple simultaneous mTurquoise2, mNeonGreen, and mScarlet-I fusions to three different proteins in single living *S. salivarius* cells. Besides FtsZ and HlpA, we selected DivIVA, as it partially co-localizes with FtsZ in *S. pneumoniae* [44]. DivIVA is a membrane-tethered hub in Gram-positive bacteria that accumulates at the septum, as well as at both cell poles due to its high affinity for negatively curved membranes [45, 46].

First, we individually evaluated DivIVA localization from the different FP fusions, because previous results in *S. pyogenes* or *Streptomyces coelicolor* indicated that DivIVA incorrectly localizes upon overexpression provoking abnormal cell morphologies [47, 48]. We first engineered constitutively expressed (P32) fusions integrated into the chromosome at the ectopic *tRNA^Ser^* locus (Supplementary Figure S4a). Akin to its profile in *S. pneumoniae* [49, 50], *S. salivarius* DivIVA localized to the midcell, and in a few cases to the cell poles (Supplementary Figure S4b). When integrated at the *tRNA^Ser^* locus and expressed under the control of two xylose-inducible promoters (P*xyl1* and P*xyl2*) [32] (Supplementary Figure S4c), we observed fluorescent DivIVA at the equatorial planes and at cell poles in *S. salivarius* cells (Supplementary Figure S4d). Expression from P*xyl1* resulted in higher fluorescence compared to P*xyl2* but morphological defects were not noticeable.

From this, we selected a combination of FtsZ-mNeonGreen, HlpA^Sp^-mTurquoise2 and DivIVA-mScarlet-I fusions in a unique triple label strain. To concomitantly observe the profiles of three distinct fusion proteins, we kept the P*32*-*hlpA^Sp^-mTurquoise2* construct at the *tRNA^Ser^* locus and transferred both the P*tet-ftsZ-mNeonGreen* and P*xyl1-divIVA-mScarlet-I* constructs to other chromosomal loci with different antibiotic markers. As integration platforms, we used two previously reported permissive loci *tRNA^Thr^* [32] and *gor* [30] (Figure 6c). In a validation experiment, we observed each fluorescent fusion separately (Figure 6d). Induction of each fusion produced a similar localization profile compared to the fusions integrated at the *tRNA^Ser^*locus (Figure 6b and S5d). Therefore, we engineered a chassis that encodes the FtsZ, DivIVA and HlpA fusions (strain VL5940: P32-*hlpA^Sp^-mTurquoise2*, P*tet-ftsZ-mNeonGreen*, P*xyl1-divIVA-mScarlet-I*) and performed a time-lapse experiment to monitor protein localization over time intracellularly (Figure 7a). Western blotting showed that the fusion proteins were correctly produced, either constitutively (HlpA^Sp^-mTurquoise2), or specifically produced upon induction with aTc and xylose for FtsZ-mNeonGreen and DivIVA-mScarlet-I, respectively (Figure 7b and Supplementary Figure S5). As expected, HlpA formed nucleoid-like condensed clusters excluded from the constriction site at late stages of division, while FtsZ and DivIVA moved from the midcell to the next area of septal formation (Figure 7c; Supplementary Movie S1). However, the timing of relocation differed between FtsZ and DivIVA, similar to the temporal hierarchy observed in *S. pneumoniae* [51]. Indeed, DivIVA colonized the midcell with a small delay and dwelled at this location for a longer time compared to FtsZ. All together, these data show that co-visualization of multiple proteins with different FPs at single cell level can be achieved in *S. salivarius*. The specific protein labeling here allowed us to refine the sequential recruitment steps of its cell division proteins in spatial context.

**Figure 7.**
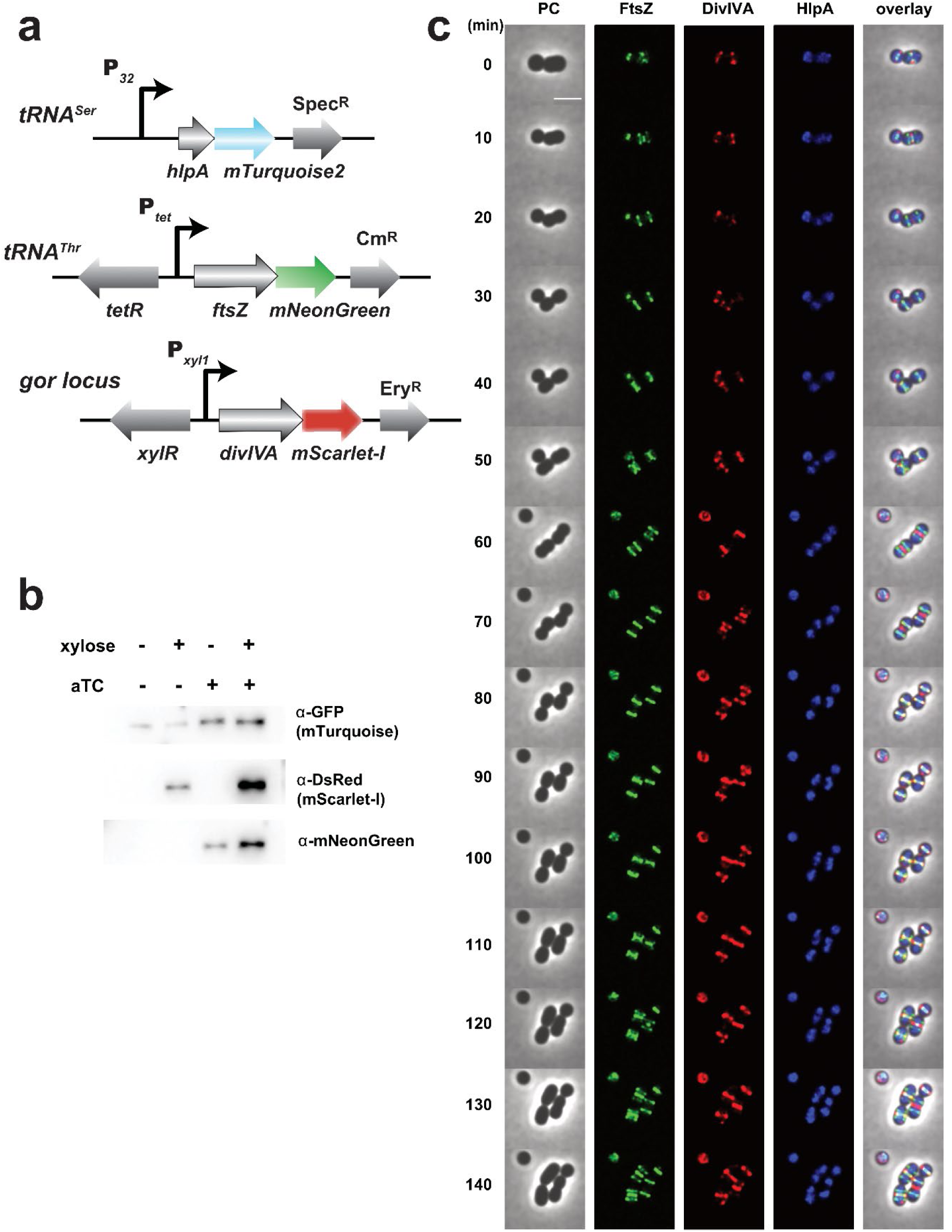
Dynamic localization of FtsZ-, DivIVA- and HlpA-fusions in *S. salivarius* cells by coexpressed fluorescent proteins. (a) Genetic constructions of the multi-labeled *S. salivarius* strain. *hlpA-mTurquoise2* was integrated at the *tRNA^Ser^* locus under the control of the constitutive P*32* promoter, while the P*tet*-inducible *ftsZ-mNeonGreen* and the P*xyl1*-inducible *divIVA-mScarlet-I* were integrated at the *tRNA^Thr^* and *gor* locus, respectively. (b) Immunoblot showing the HlpA-mTurquoise2, FtsZ-mNeonGreen, and DivIVA-mScarlet-I expression after growth in M17G with (+) or without (-) inducers (aTc and/or xylose). The mTurquoise2, mScarlet-I and mNeonGreen fusions were detected with an anti-GFP, anti-DsRed and anti-mNeonGreen primary antibodies, respectively. (c) Time-lapse images at 10 min intervals of the multi-labeled strain HlpA^Sp^-mTurquoise2 FtsZ-mNeonGreen, DivIVA-mScarlet-I, after incubation with aTc and xylose for 2 hours before imaging. The inducers were supplemented in the agarose pad used to support the cells for imaging. Fluorescent images were deconvoluted with the Huygens software and assembled with Fiji. The scale bar equals 2 μm.

## CONCLUSION

This study was based on the fluorescent labeling strategy proposed by Kjos et al. [17], which involved operon insertion of *hlpA^Sp^-fp* constructs downstream of the endogenous *hlpA^St^* gene. This approach, optimized in *S. pneumoniae*, allowed constitutive fluorescent labeling without affecting cell morphology or growth, making HlpA^Sp^-FP a reliable nucleoid, cell and population marker. Using a detailed construction protocol [8], which employs PCR amplification of five fragments, we streamlined the integration process into *S. thermophilus* and *S. salivarius* by employing a simplified three-fragment overlapping PCR strategy. Fluorescence and resistance genetic modules can be amplified from the 20 plasmids deposited at Addgene (Supplementary Table S1) to produce fluorescently-labeled bacterial strains of interest.

We validated the use of HlpA^Sp^ fused to various fluorescent proteins (mTurquoise2, mNeonGreen, msfYFP, mScarlet-I, and mKate2) and demonstrated their utility as nucleoid-localized markers across the different growth phases without detrimental effects on cell morphology or fitness in streptococci. We also showed how regulatable promoters such as the anhydrotetracycline-derepressed (P*tet*) and the xylose-inducible (P*xyl1* and P*xyl2*) promoters enabled precise temporal control of visualized protein production, minimizing the metabolic burden associated with constitutive overproduction.

Finally, we further demonstrated the potential of multi-labeling approaches, with distinct populations of HlpA-FP-labeled cells being distinguishable and allowing for simultaneous monitoring of up to three proteins or cell populations. The tools developed here provide a robust and versatile molecular toolbox, facilitating detailed studies of cellular dynamics and population behaviors in *Streptococcus* species and beyond. These findings pave the way for broader applications in microbiology, including studies involving multiple labeled populations and dynamic protein interactions.

## EXPERIMENTAL PROCEDURES

### Bacterial strains and growth conditions

Bacterial strains used in this study are listed in the Supplementary Table S3. *E. coli* was grown in LB medium with shaking at 37°C [52]. *S. thermophilus* LMG18311 and its derivatives were grown at 37°C in M17L broth (M17: Difco Laboratories Inc., MI with 1% lactose [w/v]) or in half milk [53] without shaking. *S. salivarius* HSISS4 and its derivatives were grown at 37°C in M17G broth or CDMG (M17 and CDM broth containing 1% glucose [w/v], respectively) without shaking [32]. When required, ampicillin (100 μg.ml^-1^ for *E. coli*), chloramphenicol (100 μg.ml^-1^ for *E. coli*, 4 μg.ml^-1^ for *S. thermophilus* and 5 μg.ml^-1^ for *S. salivarius*), erythromycin (150 μg.ml^-1^ for *E. coli*, 5 μg.ml^-1^ for *S. thermophilus* and 10 μg.ml^-1^ for *S. salivarius*), kanamycin (50 μg.ml^-1^ for *E. coli*, 500 μg.ml^-1^ for *S. thermophilus*), spectinomycin (50 μg.ml^-1^ for *E. coli*, 500 μg.ml^-1^ for *S. thermophilus* and 200 μg.ml^-1^ for *S. salivarius*), anhydrotetracycline (aTc; 50 ng.ml^-1^ for *S. salivarius*) or xylose (1% for *S. salivarius*) were added to the media. Synthetic peptides (95% purity) including XIP (com<u>X</u>-inducing <u>p</u>eptide for *S. thermophilus*) and sComS (for *S. salivarius*) were obtained from Peptide2.0 Inc. (Chantilly, VA, USA) and dissolved in DMSO. Plates inoculated with *S. thermophilus* or *S. salivarius* cells were incubated anaerobically (BBL GasPak systems, Becton Dickinson, NJ) at 37°C. *S. thermophilus* and *S. salivarius* transformation protocols were adapted from Dorrazehi 2020 [53] and Mignolet 2018 [32], respectively. Briefly, overnight cultures of *S. thermophilus* were diluted 1:20 and incubated for 75 min at 37°C. Then, 1 µM of XIP and 5 µL DNA cassettes (from overlapping PCRs) were added, followed by incubation for 3 h at 37°C before plating on selective M17L agar. Transformants were transferred to liquid selective M17L and after growth, re-isolated on selective solid medium. For *S. salivarius*, cultures were initiated in CDMG and incubated for 3.5 h at 37°C. Then, 1 µM sComS and DNA (from overlapping PCRs), were added, and cells were allowed to recover for 3 h at 37°C before plating on M17G agar supplemented with antibiotics when required.

### Construction of S. pneumoniae hlpA-fp

To generate *hlpA-mTurquoise2* and *hlpA-mNeonGreen* constructs in *S. pneumoniae*, the *hlpA-mTurquoise2* and *hlpA-mNeonGreen* genes were integrated downstream of the native *hlpA* gene as a second copy of *hlpA*. The up and down homology regions were amplified by PCR using the primer pairs hlpA-up-F/hlpA-up-R and cam-hlpA-down-F/cam-hlpA-down-R respectively using genomic DNA from VL1459 (*hlpA*::*hlpA*_*hlpA*-*mKate2*-Cm^R^) [54] as a template. The *mTurquoise2* and *mNeonGreen* genes were PCR-amplified with the primer pairs mTurquoise2-F/mTurquoise2-R and mNeon-F/mNeon-R, respectively. We used codon-optimized synthetic *mTurquoise2* and *mNeonGreen* genes as templates [9]. The resulting PCR fragments were fused by overlapping PCR to generate the final constructs and transformed into *S. pneumoniae* D39V.

### Construction of plasmids

Plasmids and oligonucleotides used in this study are listed in Supplementary Tables S1 and S2, respectively. Construction of *hlpA^Sp^-fp* plasmids was performed using the CloneJet PCR Cloning kit (Thermo Fisher Scientific, France) to ligate *hlpA^Sp^-fp* PCR fragments into the pJet1.2/blunt positive-selection cloning vector. Genomic DNA templates to amplify *hlpA^Sp^-fp* fragments were extracted from VL1778 (mTurquoise2), VL877 (mNeonGreen), VL1634 (msfYFP), VL1780 (mScarlet-I), or VL1459 (mKate2). The resulting recombinant plasmids contained *hlpA^Sp^-fp* inserts downstream of the T7 RNA promoter and disrupted the lethal restriction gene *eco*47I. *E. coli* strain DH5α (Invitrogen) was used as the host for plasmid amplification during cloning experiments. Electrotransformations were performed as described previously [55]. All PCR reactions were performed with the high-fidelity DNA polymerase, Phusion (ThermoScientific) according to the manufacturer’s guidelines. A fragment containing APH(3’)-IIIa (Km^R^) with its promoter was amplified from the genomic DNA of strain LMG18311kan [56] with the primer pair EcoRIKanProm_F/EcoRIK7Kanterm3. This fragment was ligated into EcoRI sites of pJet1.2 *hlpA^Sp^-mTurquoise2*/*mNeonGreen*/*msfYFP/mScarlet-I*/*mKate2* to generate plasmids with APH(3’)-IIIa located downstream of the *hlpA^Sp^-fp* genes. Fragments containing *catQ* (Cm^R^), *erm*(B) (Ery^R^) and ANT(9) (Spec^R^), along with their native promoters, were amplified from *Streptococcus salivarius* strain F4_20 (accession number LN812955.1) [57], plasmids pG^+^Host9 [58] and pSet4s [59], respectively. Primer pairs EcoRIK7catQ5’/HindK7catQ3’, EcoRIEry_F/HindEry_R or EcoRISpecFwdbis/HindSpecRev were used for the amplification. The antibiotic resistance fragments were ligated into EcoRI and HindIII sites of pJet1.2 *hlpA^Sp^-mTurquoise2*/*mNeonGreen*/*msfYFP*/*mScarlet-I*/*mKate2* to construct plasmids with *catQ*, *erm*(B) or ANT(9) located downstream of the *hlpA^Sp^-fp* genes. All plasmids were verified by sequencing and have been deposited at Addgene (plasmid #206810-206829).

### Construction of S. thermophilus hlpA^Sp^-fp -Ab^R^

DNA cassettes for *S. thermophilus* transformation were engineered using an overlapping fusion PCR procedure, as described by Mortier-Barrière *et al.* [60]. Plasmids pJet1.2 *hlpA^Sp^-mTurquoise2*/*mNeonGreen*/*msfYFP*/*mScarlet-I*/*mKate2*-Cm^R^/Ery^R^/Km^R^/Spec^R^ were used as templates for *hlpA^Sp^-fp-*Ab^R^ cassettes for PCR. All 20 different amplicons (PCR hlpA^Sp^-fp-Ab^R^) were amplified using primers hlpA-F-rbs_C, which introduce a ribosome binding site (RBS) for *hlpA^Sp^-fp* translation, and FluoAbR2. For alternative construction, the hlpA_ATG_F primer can be used to initiate at the *hlpA^Sp^* methionine start codon, and FP can be fused to the C-terminus of a protein of interest, along with Ab^R^ selection.

Homologous fragments including the *hlpA^St^* upstream and the downstream regions were amplified from strain LMG18311 (LMG strain collection) using primer pairs SthHlpANot_1/hlpA-R-rbs_B, and hlpA-down-I/SthHlpA_Apa4, respectively. The two amplicons generated, PCRhlpA_UP and PCRhlpA_DOWN, were each over 1 kb in length to promote homologous recombination. Equimolar amounts of purified PCRhlpA_UP, PCRhlpA_DOWN and PCRhlpA^Sp^-fp-Ab^R^ fragments were fused with overlapping PCRs (see Supplementary Table S4) with outers primers SthHlpANot_1 and SthHlpA_Apa4 [60]. Five microliters of the resulting amplifications were used for LMG18311 transformation and insertion via double homologous recombination downstream of the *hlpA^St^* gene. Transformants were selected on M17L supplemented with appropriate antibiotics and verified by fluorescence microscopy, PCR and sequencing.

### Construction of fp fusions in S. salivarius

The *hlpA^Sp^-fp* genes were fused with overlapping PCRs to either a strong constitutive promoter (P32) or an aTc-inducible promoter (P*tet*) and inserted with the *tetR* regulator gene (for P*tet*) and a spectinomycin resistance cassette, at the permissive tRNA serine locus (*tRNA^Ser^*; *HSISS4_r00062*) by double homologous recombination (see Supplementary Table S4). The *ftsZ-fp* genes were fused with overlapping PCRs to the P*tet* promoter and inserted by double homologous recombination at either the permissive *tRNA^Ser^* locus or the tRNA threonine locus (*tRNA^Thr^*; *HSISS4_r00061*), along with the *tetR* regulator gene and a spectinomycin or chloramphenicol resistance cassette, respectively.

The *divIVA-fp* genes were fused by overlapping PCRs to either a strong constitutive promoter (P32), or to a strong (P*xyl1*) or a mild (P*xyl2*) xylose inducible promoter and inserted by double homologous recombination at either the permissive *tRNA^Ser^* locus or the *gor* locus (downstream of *HSISS4_00325*), with the *xylR* regulator gene in the case of P*xyl1* and P*xyl2*, and with a spectinomycin or erythromycin resistance cassette, respectively. P*32*, P*xyl1* or P*xyl2* at the *tRNA^Ser^*locus were amplified from JM1101, JM1015 or JM1016, respectively [32]. The *tetR*-P*tet* fragment was amplified from VL1048. The Spec^R^ cassette at the *tRNA^Ser^* locus and the Cm^R^ cassette at the *tRNA^Thr^* locus were amplified from JM1101 and JM1100 [61], respectively. The Ery^R^ cassette was amplified from JM1029. *tRNA^Ser^*, *tRNA^Thr^*, and *gor* upstream or downstream fragments, as well *ftsZ* and *divIVA* genes, were amplified from the WT strain HSISS4. *hlpA-fp* fusions were amplified from the cognate pJet1.2 plasmid.

### Plate reader growth and fluorescence measurements

Growth curves and fluorescence were monitored using clear 96-well plates in an EnSight microplate reader (Perkin Elmer). Overnight cultures were diluted 1:100 in M17L and, after 4 hours of incubation at 37°C, adjusted to an OD590nm of 0.02 in M17L. Each sample (250 µL) was prepared in technical triplicate with the blank (medium only) and monitored at 37°C during 24h with measurement taken every 10 minutes after shaking. Absorbance was measured at a height of 7.5 mm with 50 flashes at 590 nm. Fluorescence measurements were conducted at a height of 9.5 mm with 100 flashes, using the following excitation (ex) and emission (em) settings for each fluorescent protein: 434 nm ex/474 nm em for mTurquoise2; 505 nm ex/520 nm em for mNeonGreen; 513 nm ex/530 nm em for msfYFP; 569 nm ex/594 nm em for mScarlet-I and 588 nm ex/633 nm em for mKate2.

### Sample preparation and Immunoblotting

1. *S. salivarius* cells were overnight-precultured in M17G, diluted the next morning to a final OD∼0.01 in fresh M17G supplemented with aTc and/or xylose, if required, and grown for 6h to reach mid-exponential phase. Cells were harvested by centrifugation (10 min; 4,050 x g). Supernatants were discarded, and the cell pellets were resuspended in 1 ml of cold PBS. To normalize cells quantity across samples, ODs were measured, cells were centrifuged, and pellets were resuspended in specific volumes of buffer A (50mM Tris-HCl, 50 mM NaCl, 1mM EDTA, pH 8,0), supplemented with protease inhibitors (Sigma 324890) and 100 µl Zirconia beads (Biospec, 0612731). Cell lysis was performed using a FastPrep homogenizer (MP biomedicals) for 2 cycles of 40 sec at 6 m.s^-1^, with 40 sec on ice between cycles. Lysates were centrifuged (1 min, 4°C, 13.000 x g) and 100 µl of supernatants were mixed with 20 µl of SDS loading dye. Samples were heated at 95°C for 10 min and cooled down on ice for an additional 10 min. Crude extracts were separated by SDS-PAGE and blotted on methanol-activated PVDF (polyvinylidenfluoride) membranes (Merck Millipore). Membranes were blocked for 1 h with Tris-buffered saline (20mM Tris base 150 mM NaCl), 0.05% (v/v) Tween 20 (TBS-T), and 5% (w/v) dry milk, followed by a 1-hour incubation with the primary antibodies diluted in TBS-T, containing 5% dry milk. The membranes were washed four times for 5 minutes each in TBS-T and incubated for 1 hour with the secondary antibody diluted in TBS-T with 5% dry milk. After four additional 5 min washes in TBS-T, the membranes were developed using Super signal West Pico PLUS Chemiluminescent substrate (Protein Biology, 34577) and visualized with the Fusion Fx Camera (Vilber Lourmat). Primary antibodies included rabbit antisera against mTurquoise2 (anti-GFP, Invitrogen, A6455) and mScarlet-I (anti-DsRed, Takara, 632496), as well as Mouse monoclonal antibodies against mNeonGreen (ChromoTek, 32F6), all at 1:5,000 dilution. HRP-conjugated Goat anti-rabbit (Abcam, AB205718) and anti-mouse (Promega, W402B) secondary antibodies were used at a 1:5,000 and 1:2,500 dilutions, respectively.

### Image acquisition

*S. thermophilus* cells were grown overnight in M17L diluted to an OD of 0.01 in fresh M17L. After 5 hours of incubation, 1µL of culture or mixed cultures were spotted on a 1% agarose pad. Imaging was performed using a Nikon Eclipse Ti-E inverted microscope with a perfect focus system (PFS), pE-100 CoolLED, a Plan Apo λ 100 × 1.45 oil objective (Nikon) and a Hamamatsu ORCA-Flash4.0 V2 C11440-22CU camera (Hamamatsu, Hamamatsu City, Japan). Phase-contrast images were acquired using transmitted light and an exposure time of 20 ms. Epifluorescence microscope laser and detector settings were optimized to discriminate between the different fluorescence signals. Snapshot fluorescence images were acquired with a laser intensity set at 10% and exposure times of 200 ms for mTurquoise2, 100 ms for mNeonGreen, 300 ms for msfYFP and 300 ms for mScarlet-I to minimize photobleaching. Images were exported as 16-bit TIF files. The Nikon Ti filters sets used were: CFP HQ (Ex: 436/20 nm, DM: 455 nm, BA: 480/40 nm), GFPHQ (Ex: 470/40 nm, DM: 495 nm, BA: 525/50 nm), YFPHQ (Ex: 500/20 nm, DM: 515 nm, BA: 535/50 nm), Texas Red (Ex: 560/40 nm, DM: 595 nm, BA: 630/60 nm) as described in Daveri 2023 [62].

*S. salivarius* cells were grown overnight in CDMG without any inducer and diluted to an OD of 0.01 in fresh CDMG. After 2 hours of incubation, when cultures reached an OD of ∼0.1, culture media were supplemented with aTc and/or xylose as appropriate and the incubation continued for an additional 2 hours. To image exponentially growing cultures, 1µl of culture was spotted onto PBS 1% agarose pad. For time-lapse microscopy, 0.7µl of cells were spotted onto CDMG 1% agarose pad supplemented with aTc and xylose. The pad was maintained at 30°C for the duration of imaging. Pads were placed inside a gene frame (Thermo Fisher Scientific) and sealed with a cover glass as described previously [42]. Microscopy was conducted using a Leica DMi8 microscope equipped with a sCMOS DFC9000 GT (Leica) camera and a SOLA light engine (Lumencor’s SPECTRA; 7-channel, solid-state light source for epi-fluorescence imaging) and a ×100/1.40 oil-immersion objective. Phase-contrast images were acquired using transmission light with a 50 ms exposure. Snapshot fluorescence images were acquired with a 700 ms exposure, while time-lapse laser intensity (50%) and exposure times were optimized to minimize photobleaching with an exposure time of 300 ms for mTurquoise2, 400 ms for mNeonGreen, and 1000 ms for mScarlet-I. The filter sets used included: mTurquoise2 (Ex: 440 nm, BS: 455 nm Chroma, Em: 470/26 nm Chroma ET470/26 m), mNeonGreen (Ex: 490/20 nm, BS: 510 nm, Em: 525/36 nm Chroma ET535/30 m), msfYFP (Ex: 510 nm, BS: 520 nm Chroma 69008, Em: 535/30 nm Chroma ET535/30 m), mScarlet-I (Ex: 550 nm Chroma ET545/30 X, BS: 595 nm Chroma 69008, Em: 635/70 nm Chroma ET635/70 m), and mKate2 (Ex: 550 nm Chroma ET575/30, BS: 595 nm Chroma 69008, Em: 635/70 nm Chroma ET635/70 m). Images acquisition was performed using LasX v.3.4.2.18368 software (Leica).

### Image processing

All microscope images were processed with FIJI v2.14.0 to adjust contrast and brightness. Adjustments were automated to maintain consistent settings across all images within a single panel. Signal-to-noise ratio quantification was performed by segmentation on three independent images for each HlpA-FP fusion. The mean background (noise) value and the mean value intensity (signal) values within segmented cells were extracted for analysis. Deconvolution of time-lapse images was performed with the Huygens v.17.10.0p4 (SVI) software. Default settings were applied, and the number of iterations was optimized for mTurquoise2 (9x), mNeonGreen (9x), and mScarlet-I (12x). Time-lapse images were corrected for drift to mount the time-lapse movie.

## SUPPLEMENTAL INFORMATION

Supplemental Information includes six supplementary figures, one supplementary movie, and four supplementary tables.

## AUTHORS CONTRIBUTION

J.M., J.R.vdM., J.-W.V., and V.L. Conception and design. J.M., J.S. and V.L. Acquisition of data. J.M., J.R.vdM., J.-W.V., and V.L. Analysis and interpretation of data. J.M., J.R.vdM., J.-W.V., and V.L. Draft or revising the article.

## ACKNOWLEDGEMENTS

We are extremely grateful to Profs. Mortens Kjos and Jun Kurushima for the *S. pneumoniae hlpA*-*fp* strains. The work of J.-W.V. was supported by the Swiss National Science Foundation (SNSF grants 310030_192517, 310030_200792 and NCCR 51NF40_180541). J.M. received funding from the European Union’s Horizon 2020 research and innovation program (Marie Skłodowska-Curie grant N°101018461). The work of V.L. was supported by the Université de Lorraine, Agence Nationale de la Recherche (ANR) throught the LUE Widen Horizons grant and by the French National Research Institute for Agriculture Food and Environment (INRAE).

## COMPETING INTERESTS

The authors declare they have no conflict of interests.

## Notes

### Competing Interest Statement

The authors have declared no competing interest.

