## Supplementary material for "A Modular Genetic Toolbox for Precise Gene Regulation and Multi-Color Imaging in Streptococci": Figure S1 to S5

### Supplemental Information

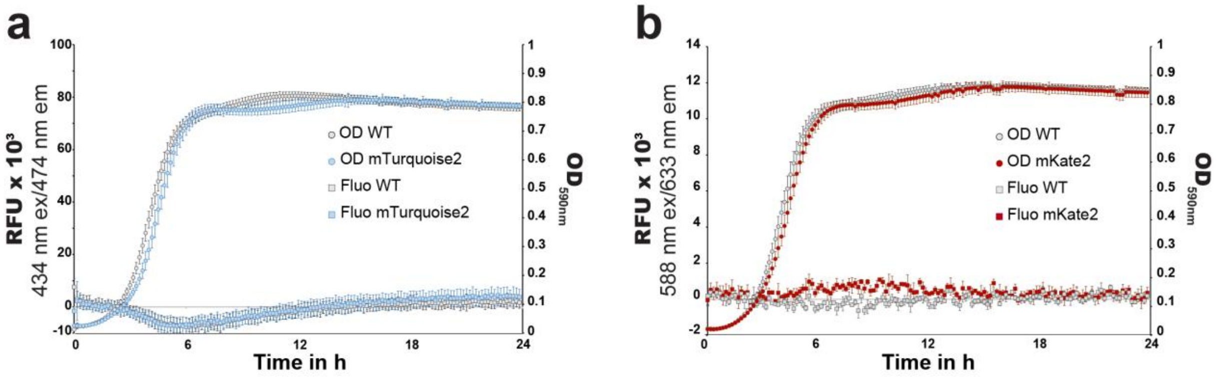


**Supplementary Figure S1. Growth curves and fluorescence quantifications of *S. thermophilus hlpA$^{Sp}$-mTurquoise2* and *hlpA$^{Sp}$-mKate2* strains.**

Optical density ($OD_{590nm}$, round symbols) and fluorescence measurements, expressed in Relative Fluorescence Units (RFUs, square symbols), of *S. thermophilus* LMG18311 and its derivative strains grown in M17L medium without antibiotics over 24 hours post-dilution. Each LMG18311 derivative is represented by its distinct colored plot symbol, while the absence of fluorescence in the parental LMG18311 strain is plotted in grey. RFU measurements were taken at 434 nm ex/474 nm em for mTurquoise2 (**a**) and 588 nm ex/633 nm em for mKate2 (**b**). All experiments were performed in triplicate.

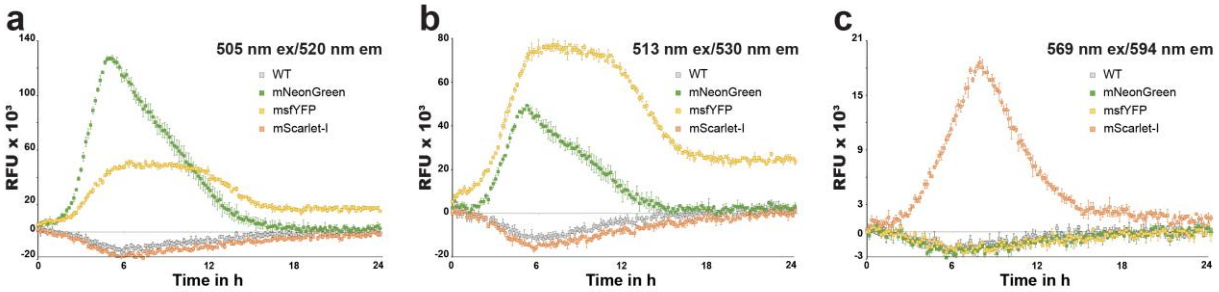


**Supplementary Figure S2. Signal bleed-through for HlpA$^{Sp}$-FP in *S. thermophilus*.**

Comparative RFU values of the 3 fluorescent protein fusions HlpA$^{Sp}$-mNeonGreen, HlpA$^{Sp}$-msfYFP and HlpA$^{Sp}$-mScarlet-I measured at each of their specific excitation/emission wavelengths. Each engineered LMG18311 strain is represented by its distinct colored square symbol, while the absence of fluorescence of the parental LMG18311 strain is plotted in grey. RFU measurements were taken at 505 nm ex/520 nm em (**a**), 513 nm ex/530 nm em (**b**), and 569 nm ex/594 nm em (**c**). All experiments were performed in triplicate.

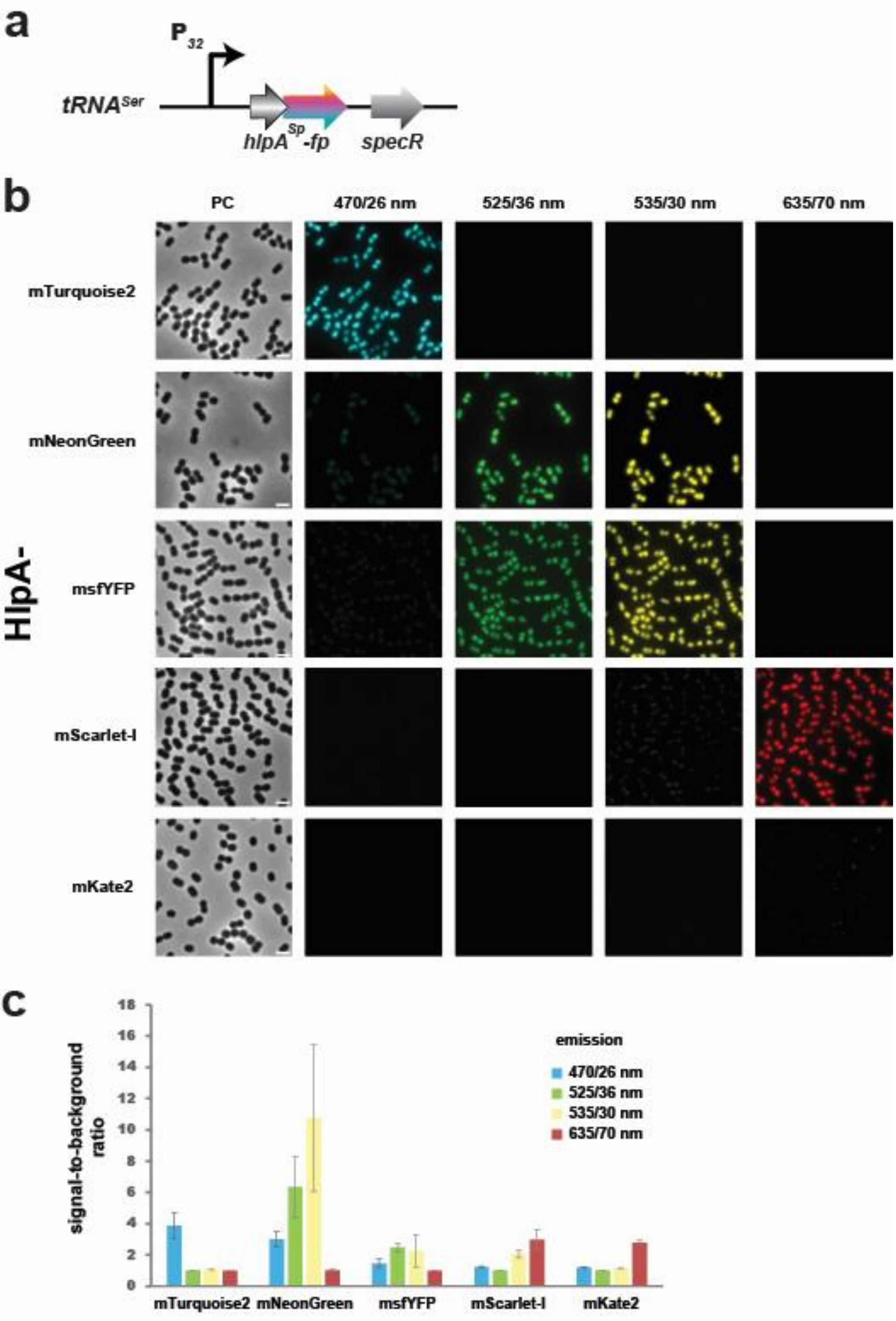
a
P32
tRNASer
hlpASp-fp
specR
b
PC
470/26 nm
525/36 nm
535/30 nm
635/70 nm
HlpA-
mTurquoise2
mNeonGreen
msfYFP
mScarlet-I
mKate2
c
signal-to-background ratio
emission
470/26 nm
525/36 nm
535/30 nm
635/70 nm
mTurquoise2
mNeonGreen
msfYFP
mScarlet-I
mKate2

21

**Supplementary Figure S3. Fluorescence channel specificity for 5 FPs in *S. salivarius***

(a) Scheme of the genomic topology of the constitutive $P_{32}$ controlling the *hlpA-fp* integrated at the $tRNA^{Ser}$ locus. The $Spec^R$ gene is used to select chromosomal integration of *hlpA-fp* constructs.

(b) Fluorescent protein tags of the $HlpA^{Sp}$ nucleoid-associated factor were observed under microscope for single-cell analyses. Phase contrast (PC) and fluorescence recorded in the four channels to assess bleed-through are displayed for mTurquoise2, mNeonGreen, msfYFP, mScarlet-I and mKate2 fluorescent tags. The same brightness/contrast parameters were applied to the fluorescence images to evaluate the bleed-through. The scale bar equals 2 µm.

(c) Quantification histogram of signal-to-noise ratio for five FPs in the 4 cognate emission channels. Images were segmented and analyzed with the Fiji software to elicit the mean fluorescence intensity per cell. Noise was estimated as the mean value of the background (image area excluding the cell bodies). Experimental values represent the averages (with SEM) of three independent images.

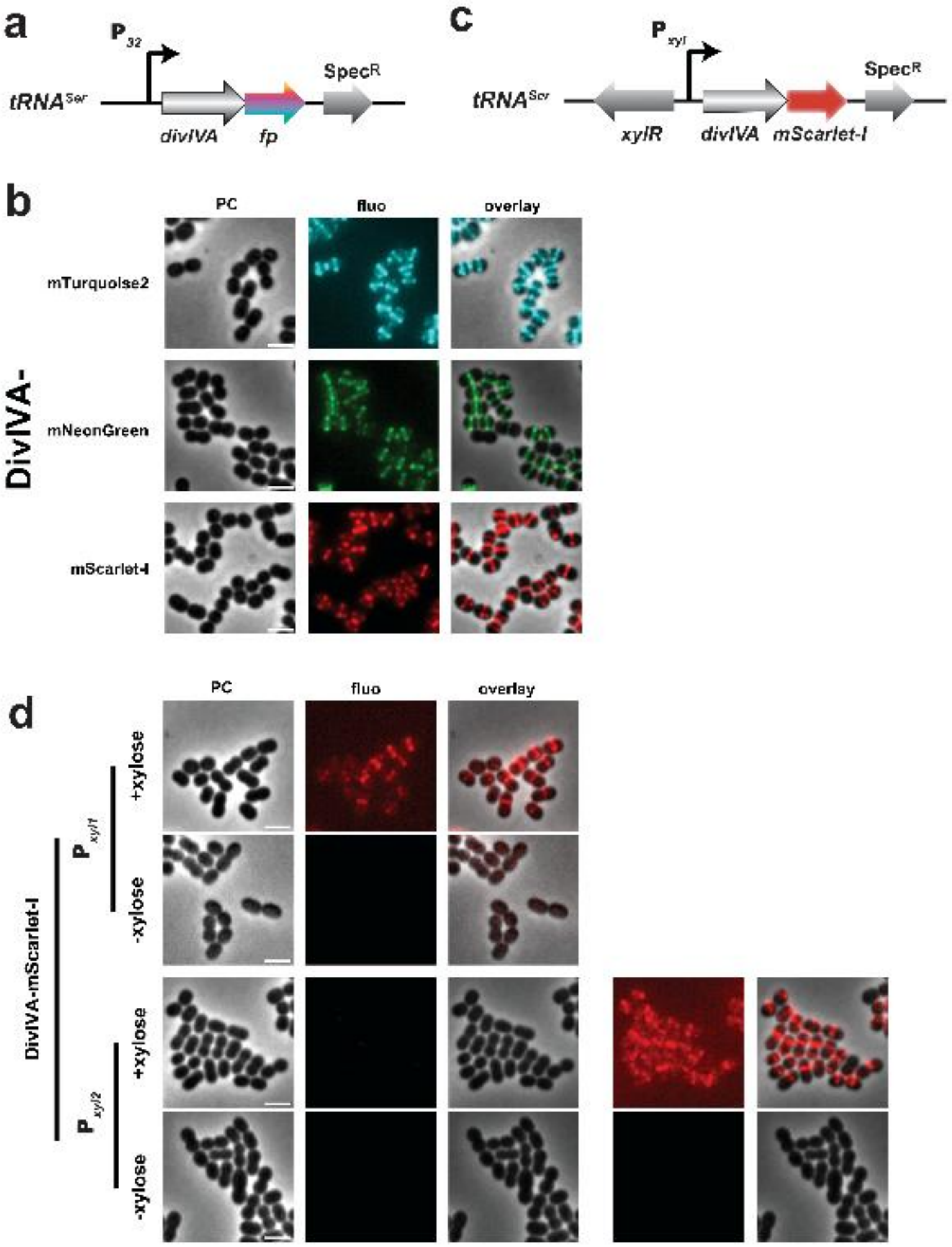


**Supplementary Figure S4. DivIVA subcellular localization in *S. salivarius***

(a) Scheme of the genomic topology of the constitutive $P_{32}$ controlling the *divIVA-fp* integrated at the *tRNA*$^{Ser}$ locus. The Spec$^{R}$ gene is used to select chromosomal integration of *divIVA-fp* constructs.

(b) Single-cell imaging of constitutively-produced tagged DivIVA. mTurquoise2, mNeonGreen and mScarlet-I fusion mutants were imaged in mid-exponential phase. Phase contrast (PC), fluorescence recorded in the cognate channel, and overlay images are depicted. The scale bar equals 2 µm.

(c) Scheme of the genomic topology of the $P_{xyl}$-inducible *divIVA-mScarlet-I* integrated at the *tRNA*$^{Ser}$ locus. The *xylR* repressor genes, with its own promoter (not depicted here), points in the opposite orientation compared to the strong $P_{xyl1}$ or mild $P_{xyl2}$ inducible promoter. The Spec$^{R}$ gene is used to select chromosomal integration of *divIVA-fp* constructs.

(d) Single-cell imaging of xylose-inducible tagged DivIVA. mScarlet-I fusion mutants were incubated with xylose for 2 hours before imaging. Phase contrast (PC), fluorescence, and overlay images are depicted. The same brightness/contrast parameters were applied to the fluorescence images to qualitatively compare the intensities between the $P_{xyl1}$ or $P_{xyl2}$ expression system (left panel). On the bottom right, the brightness/contrast parameters were adjusted to visualize the fluorescent signal in the $P_{xyl2}$ expression system. The scale bar equals 2 µm.

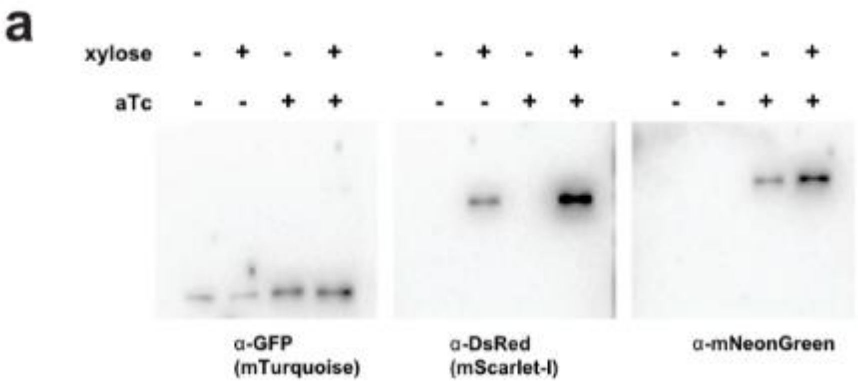


**Supplementary Figure S5.**

Full-scan image of immunoblot showing the HlpA-mTurquoise2, FtsZ-mNeonGreen, and DivIVA-mScarlet-I steady-state levels. Cells were grown in M17G supplemented with (+) or without (-) inducing molecules (aTc and/or xylose). The mTurquoise2, mScarlet-I and mNeonGreen fusions were detected with an anti-GFP, anti-DsRed and anti-mNeonGreen primary antibodies, respectively.

**Supplementary Movie S1**

Time-lapse movie of the multi-labeled strain $HlpA^{Sp}$-mTurquoise2 FtsZ-mNeonGreen DivIVA-mScarlet-I. Cells were incubated with aTc and xylose for 2 hours before imaging. The inducing molecules were supplemented in the agarose pad. Phase contrast (PC), fluorescence recorded in the three cognate channels, and overlay images are depicted. Individual fluorescent images were deconvoluted with the Huygens software and assembled with Fiji. The scale bar equals 2 µm.

**Supplementary Table S1 Plasmids**

| | Characteristics | Reference / #plasmid |
|---|---|---|
| pJet1.2 | blunt positive-selection cloning vector *bla* | Invitrogen |
| pJethlpAmKate2Cm | pJet1.2 *hlpA*$^{Sp}$-*mKate2 catQ* | This work / #206810 |
| pJethlpAmKate2Ery | pJet1.2 *hlpA*$^{Sp}$-*mKate2 erm*(B) | This work / #206811 |
| pJethlpAmKate2Km | pJet1.2 *hlpA*$^{Sp}$-*mKate2* APH(3’)-IIIa | This work / #206812 |
| pJethlpAmKate2Spc | pJet1.2 *hlpA*$^{Sp}$-*mKate2* ANT(9*)* | This work / #206813 |
| pJethlpAmNeonGreenCm | pJet1.2 *hlpA*$^{Sp}$-*mNeonGreen catQ* | This work / #206814 |
| pJethlpAmNeonGreenEry | pJet1.2 *hlpA*$^{Sp}$-*mNeonGreen erm*(B) | This work / #206815 |
| pJethlpAmNeonGreenKm | pJet1.2 *hlpA*$^{Sp}$-*mNeonGreen* APH(3’)-IIIa | This work / #206816 |
| pJethlpAmNeonGreenSpc | pJet1.2 *hlpA*$^{Sp}$-*mNeonGreen* ANT(9) | This work / #206817 |
| pJethlpAmScarletCm | pJet1.2 *hlpA*$^{Sp}$-*mScarlet-I catQ* | This work / #206818 |
| pJethlpAmScarletEry | pJet1.2 *hlpA*$^{Sp}$-*mScarlet-I erm*(B) | This work / #206819 |
| pJethlpAmScarletKm | pJet1.2 *hlpA*$^{Sp}$-*mScarlet-I* APH(3’)-IIIa | This work / #206820 |
| pJethlpAmScarletSpc | pJet1.2 *hlpA*$^{Sp}$-*mScarlet-I* ANT(9) | This work / #206821 |
| pJethlpAmTurquoise2Cm | pJet1.2 *hlpA*$^{Sp}$-*mTurquoise2 catQ* | This work / #206822 |
| pJethlpAmTurquoise2Ery | pJet1.2 *hlpA*$^{Sp}$-*mTurquoise2 erm*(B) | This work / #206823 |
| pJethlpAmTurquoise2Km | pJet1.2 *hlpA*$^{Sp}$-*mTurquoise2* APH(3’)-IIIa | This work / #206824 |
| pJethlpAmTurquoise2Spc | pJet1.2 *hlpA*$^{Sp}$-*mTurquoise2* ANT(9) | This work / #206825 |
| pJethlpAYFPCm | pJet1.2 *hlpA*$^{Sp}$-*msfYFP catQ* | This work / #206826 |
| pJethlpAYFPEry | pJet1.2 *hlpA*$^{Sp}$-*msfYFP erm*(B) | This work / #206827 |
| pJethlpAYFPKm | pJet1.2 *hlpA*$^{Sp}$-*msfYFP* APH(3’)-IIIa | This work / #206828 |
| pJethlpAYFPSpc | pJet1.2 *hlpA*$^{Sp}$-*msfYFP* ANT(9) | This work / #206829 |

**Supplementary Table S2 Oligonucleotides**

| Names | Sequences |
|---|---|
| hlpA-up-F | 5’-AACAAGTCAGCCACCTGTAG-3’ |
| hlpA-up-R | 5’-TGATCCTTTAGCTGCAGCTTCTCCACC-3’ |
| cam-hlpA-down-F | 5’-TAAGAATTCTAATGAGCACTAGTAGG-3’ |
| cam-hlpA-down-R | 5’-CGTGGCTGACGATAATGAGG-3’ |
| mTurquoise2-F | 5’-AAGCTGCAGCTAAAGGATCAGTGAGCAAGGGCGAGGAGCTG-3’ |
| mTurquoise2-R | 5’-AGTGCTCATTAGAATTCTTATTACTTGTACAGCTCGTCCATGCCGAG-3’ |
| mNeon-F | 5’-AAGCTGCAGCTAAAGGATCAGTCTCTAAAGGTGAAGAAGATAATATGG-3’ |
| mNeon-R | 5’-AGTGCTCATTAGAATTCTTATTATTTATACAATTCATCCATACCC-3’ |
| hlpA_LB_F | 5’-CATGGCAAACAAACAAGATTTGATCGCTAAAG-3’ |
| Cm-int-R | 5’-AAAGTCGTTTGTTGGTTCAAATAATGAT-3’ |
| EcoRIKanProm_F | 5’-CCGAATTCACGTTAACAACCGGTACCTC-3’ |
| EcoRIK7Kanterm3 | 5’-CGGGAAATTTGCAGAGCAGGTAGCTATTC-3’ |
| EcoRIK7catQ5' | 5’-CCGAATTCCGTATTGGTGGAATAACACG-3’ |
| HindK7catQ3' | 5’-CTGAAGCTTGCTCTGCTTAGGATAAATAGC-3’ |
| EcoRIEry_F | 5’-CCGAATTCCAAACTTAAGAGTGTGTTGATAGTGC-3’ |
| HindEry_R | 5’-CTGAAGCTTGGACCTCTTTAGCTCCTTGG-3’ |
| EcoRISpecFwdbis | 5’-CCGAATTCCGAAGTTATCGTAACGTGAC-3’ |
| HindSpecRev | 5’-CTGAAGCTTCCAATTAGAATGAATATTTCCC-3’ |
| hlpA-F-rbs_C | 5’-GCATGCTGGAGGAATCATTAACATGGCAAACAAACAAGATTTGA-3’ |
| hlpA_ATG_F | 5’-ATGGCAAACAAACAAGATTTG-3’ |
| FluoAbR2 | 5’-TGGATGCTATTAACCCTGAACTTTCTT-3’ |
| SthHlpANot_1 | 5’-GGGCGGCCGCGTCAGGTGAGTAAGGTATG-3’ |
| hlpA-R-rbs_B | 5’-GTTAATGATTCCTCCAGCATGCGGGATTAACATTATTTGACTGCGT-3’ |
| hlpA-down-I | 5’-GTTCAGGGTTAATAGCATCCACCTAAGCTTTTAGGGGTTTTGA-3’ |
| SthHlpA_Apa4 | 5’-CGGGGCCCGTCAGAAGCTTATACCTTATCC-3’ |
| UF_tRNAser | 5’-CAAGATTAACCATGACCTTC-3’ |
| DR_tRNAser2 | 5’-TTGGATAAGGTCTTGACTTC-3’ |
| R_P32 | 5’-CATCTAAATTCCTCCTCTAG-3’ |
| F_spec | 5’-TAATAAGGCCGGCCAATAAA-3’ |
| F_hlpA_(P32) | 5’-CTAGAGGAGGAATTTAGATGGCAAACAAACAAGATTTGA-3’ |
| R_mKate2 | 5’-TTTATTGGCCGGCCTTATTAACGGTGTCCCAATTTAC-3’ |
| R_neonGreen | 5’-TTTATTGGCCGGCCTTATTATTTATACAATTCATCC-3’ |
| R_Scarlet | 5’-TTTATTGGCCGGCCTTATTATTTATATAGTTCGTCC-3’ |
| R_mTurquoise | 5’-TTTATTGGCCGGCCTTATTACTTGTACAGCTCGTCC-3’ |
| R_YFP | 5’-TTTATTGGCCGGCCTTATTATTTATAAAGTTCGTCC-3’ |
| UR_tRNAser | 5’-AGTAATTAAAAAGAAGATGG-3’ |
| F_tetR_(ser) | 5’-CCATCTTCTTTTTAATTACTCTATTCATTGTTTTGCATGC-3’ |
| R_Ptet | 5’-CATTATTTTTCCTCCTTATTT-3’ |
| F_hlpA_(Ptet) | 5’-AAATAAGGAGGAAAAATAATGGCAAACAAACAAGATTTGA-3’ |
| F_linker-fp | 5’-AGAGGATCTGGTGGAGAAG-3’ |
| F_ftsZ_(Ptet) | 5’-AAATAAGGAGGAAAAATAATGAGTTTTTCATTTGATAGC-3’ |
| R_ftsZ_linkerfp | 5’-CTTCTCCACCAGATCCTCTACGATTTTTAAAGAATGGTG-3’ |
| F_divIVA_(P32) | 5’-CTAGAGGAGGAATTTAGATGGCTATTACAGCACTTG-3’ |
| R_divIVA_linkerfp | 5’-CTTCTCCACCAGATCCTCTTTCGCTAATATTCAATTTAAAC-3’ |
| R_pZX9_ATG | 5’-CATATTTACCTCCTTTGATTTA-3’ |
| F_DivIVA_(Pxyl) | 5’-TAAATCAAAGGAGGTAAATATGGCTATTACAGCACTTG-3’ |
| UF_tRNAthr | 5’-TGTCAAAGGATTAGGAAAAC-3’ |
| UR_tRNAthr | 5’-TTGATTTATACCTCTCAATTT-3’ |
| F_cat | 5’-TAGACCCCGGGGATCCTC-3’ |
| DR_tRNAthr | 5’-AAAAAAGAATTCATTCATGATGAGCGGGTTCGTGAGA-3’ |
| F_tetR_(thr) | 5’-AAATTGAGAGGTATAAATCAACTATTCATTGTTTTGCATGC-3’ |
| R_nGreen_(cat) | 5’-GAGGATCCCCGGGGTCTATTATTTATACAATTCATCC-3’ |

| | |
|---|---|
| Fw.Up.gor | 5’-GGTGTTAAATTGACTGAAAAAG-3’ |
| Rev.Up.gor | 5’-TGAGTGAATGGTTTCAATTG-3’ |
| DF_GOR_(ery) | 5’- TACATTCCCTTTAGTAACGTGAAAACGGGTTCAGAAGAATTTG-3’ |
| Rev.Dn.gor | 5’-GCTCAAACATTTTCTAAGATTAC-3’ |
| F_xylR_GOR | 5’-CAATTGAAACCATTCACTCATCTAGATTATATATGATATGATC-3’ |
| R_fp_(ery) | 5’-CCTTATGGGATTTATCTTCCTTAAAGAATCTTGCTTGGCAAGG-3’ |
| Uplox66 | 5’-TAAGGAAGATAAATCCCATAAGG-3’ |
| DNlox71 | 5’-TTCACGTTACTAAAGGGAATGTA-3’ |

**Supplementary Table S3 Strains**

| | Characteristics | Reference/source |
|---|---|---|
| ***Escherichia coli*** | | |
| DH5α | *supE hsd-5 thi (lac-proAB) F (traD6 proAB lacIq lacZ M15) repA, derivative of strain TG1 (56) repA, derivative of strain JM101* | Invitrogen |
| ***Streptococcus thermophilus*** | | |
| LMG18311 | Wild-type strain | BCCM/LMG, strain collection |
| SC00501 | LMG18311 *hlpA-mTurquoise2*-Ery$^R$ | This work |
| SC00502 | LMG18311 *hlpA-mNeonGreen*-Ery$^R$ | This work |
| SC00503 | LMG18311 *hlpA-msfYFP*-Ery$^R$ | This work |
| SC00504 | LMG18311 *hlpA-mScarlet-I*-Ery$^R$ | This work |
| SC00505 | LMG18311 *hlpA-mKate2*-Ery$^R$ | This work |
| ***Streptococcus salivarius*** | | |
| HSISS4 | Wild-type gastro-intestinal tract isolate | (Van den Bogert et al. 2014) |
| JM1101 | HSISS4 *tRNA*$^{Ser}$::P$_{32}$-*scuR*-Spec$^R$ | (Mignolet et al. 2019) |
| JM1015 | HSISS4 *tRNA*$^{Ser}$::P$_{xyl1}$-*comR*-Spec$^R$ | (Mignolet et al. 2018) |
| JM1016 | HSISS4 *tRNA*$^{Ser}$::P$_{xyl2}$-*comR*-Spec$^R$ | (Mignolet et al. 2018) |
| JM1100 | HSISS4 *tRNA*$^{Thr}$::P$_{sptA}$-*luxAB*-Cm$^R$ | (Mignolet et al. 2019) |
| JM1029 | HSISS4 *tRNA*$^{Thr}$::P$_{comS}$-*luxAB*-Cm$^R$ Δ*comRS*::Ery$^R$ | (Mignolet et al. 2018) |
| VL5850 | HSISS4 *tRNA*$^{Ser}$::P$_{32}$-*hlpA-mNeonGreen*-Spec$^R$ | This work |
| VL5851 | HSISS4 *tRNA*$^{Ser}$::P$_{32}$-*hlpA-mScarlet-I*-Spec$^R$ | This work |
| VL5852 | HSISS4 *tRNA*$^{Ser}$::P$_{32}$-*hlpA-mTurquoise2*-Spec$^R$ | This work |
| VL5942 | HSISS4 *tRNA*$^{Ser}$::P$_{32}$-*hlpA-mKate2*-Spec$^R$ | This work |
| VL5943 | HSISS4 *tRNA*$^{Ser}$::P$_{32}$-*hlpA-msfYFP*-Spec$^R$ | This work |
| VL5853 | HSISS4 *tRNA*$^{Ser}$::P$_{tet}$-*hlpA-mNeonGreen*-Spec$^R$ | This work |
| VL5854 | HSISS4 *tRNA*$^{Ser}$::P$_{tet}$-*hlpA-mScarlet-I*-Spec$^R$ | This work |
| VL5855 | HSISS4 *tRNA*$^{Ser}$::P$_{tet}$-*hlpA-mTurquoise2*-Spec$^R$ | This work |
| VL5856 | HSISS4 *tRNA*$^{Ser}$::P$_{tet}$-*ftsZ-mNeonGreen*-Spec$^R$ | This work |
| VL5857 | HSISS4 *tRNA*$^{Ser}$::P$_{tet}$- *ftsZ -mScarlet-I*-Spec$^R$ | This work |
| VL5858 | HSISS4 *tRNA*$^{Ser}$::P$_{tet}$- *ftsZ -mTurquoise2*-Spec$^R$ | This work |
| VL5859 | HSISS4 *tRNA*$^{Ser}$::P$_{32}$-*divIVA-mNeonGreen*-Spec$^R$ | This work |
| VL5860 | HSISS4 *tRNA*$^{Ser}$::P$_{32}$- *divIVA -mScarlet-I*-Spec$^R$ | This work |
| VL5861 | HSISS4 *tRNA*$^{Ser}$::P$_{32}$- *divIVA -mTurquoise2*-Spec$^R$ | This work |
| VL5894 | HSISS4 *tRNA*$^{Ser}$::P$_{xyl1}$- *divIVA -mScarlet-I*-Spec$^R$ | This work |
| VL5895 | HSISS4 *tRNA*$^{Ser}$::P$_{xyl2}$- *divIVA -mScarlet-I*-Spec$^R$ | This work |
| VL5930 | HSISS4 *tRNA*$^{Thr}$::P$_{tet}$-*ftsZ-mNeonGreen*-Cm$^R$ | This work |
| VL5932 | HSISS4 *gor*::P$_{xyl1}$- *divIVA -mScarlet-I*-Ery$^R$ | This work |
| VL5931 | VL5852 *tRNA*$^{Thr}$::P$_{tet}$-*ftsZ-mNeonGreen*-Cm$^R$ | This work |
| VL5940 | VL5931 *gor*::P$_{xyl1}$- *divIVA -mScarlet-I*-Ery$^R$ | This work |
| ***Streptococcus pneumoniae*** | | |
| VL1 | D39V | (Slager et al. 2018) |
| VL1048 | D39V p*PEP*:: *P*$_{F6}$-*tetR*-*P*$_{T5\text{-}3}$-*luc-gfp* | (Sorg et al. 2020) |
| VL877 | D39V *hlpA*:: *hlpA_hlpA-mNeonGreen*-Cm$^R$ | This work |
| VL1459 | D39V *hlpA*:: *hlpA_hlpA-mKate2*-Cm$^R$ | (Kjos and Veening 2014) |
| VL1634 | D39V *hlpA*:: *hlpA_hlpA-msfYFP*-Cm$^R$ | Lab collection, unpublished |
| VL1778 | D39V *hlpA*:: *hlpA_hlpA-mTurquoise2*-Cm$^R$ | This work |
| VL1780 | D39V *hlpA*:: *hlpA_hlpA-mScarlet-I*-Cm$^R$ | (Kurushima et al. 2020) |

**Supplementary Table S4 Overlapping and cloning PCR Subfragments**

| PCR | Primer 1 | Primer 2 |
|---|---|---|
| Upstream homologous region of *hlpA* in *S. pneumoniae* | hlpA-up-F | hlpA-up-R |
| Downstream homologous region of *hlpA* in *S. pneumoniae* | cam-hlpA-down-F | cam-hlpA-down-R |
| *mTurquoise2* amplification | mTurquoise2-F | mTurquoise2-R |
| *mNeonGreen* amplification | mNeon-F | mNeon-R |
| Km$^R$ cassette | EcoRIKanProm_F | EcoRIK7Kanterm3 |
| Cm$^R$ cassette | EcoRIK7catQ5' | HindK7catQ3' |
| Ery$^R$ cassette | EcoRIEry_F | HindEry_R |
| Spec$^R$ cassette | EcoRISpecFwdbis | HindSpecRev |
| $hlpA^{Sp}$-*fp* amplifications to clone in pJet1.2 | hlpA_LB_F | Cm-int-R |
| $hlpA^{Sp}$-*fp*-Ab$^R$ amplifications to introduce a RBS (PCRhlpASp-fp-Ab$^R$) | hlpA-F-rbs_C | FluoAbR2 |
| $hlpA^{Sp}$-*fp*-Ab$^R$ amplifications | hlpA_ATG_F | FluoAbR2 |
| Upstream homologous region (including $hlpA^{St}$) of $hlpA^{St}$ locus (PCRhlpA_UP) | SthHlpANot_1 | hlpA-R-rbs_B |
| Downstream homologous region of $hlpA^{St}$ locus (PCRhlpA_DOWN) | hlpA-down-I | SthHlpA_Apa4 |
| Upstream homologous region of $tRNA^{Ser}$ locus | UF_tRNAser | UR_tRNAser |
| Upstream homologous region of $tRNA^{Thr}$ locus | UF_tRNAthr | UR_tRNAthr |
| Upstream homologous region of *gor* locus | Fw.Up.gor | Rev.Up.gor |
| Downstream homologous region of *gor* locus (for $ery^R$ fusion) | DF_GOR_(ery) | Rev.Dn.gor |
| Ery$^R$ cassette amplification | Uplox66 | DNlox71 |
| Spec$^R$ cassette at the $tRNA^{Ser}$ locus | F_spec | DR_tRNAser2 |
| Cm$^R$ cassette at the $tRNA^{Thr}$ locus | F_cat | DR_tRNAthr |
| $P_{32}$ at the $tRNA^{Ser}$ locus | UF_tRNAser | R_P32 |
| $P_{xyl1}$ at the $tRNA^{Ser}$ locus | UF_tRNAser | R_pZX9_ATG |
| $P_{xyl2}$ at the $tRNA^{Ser}$ locus | UF_tRNAser | R_pZX9_ATG |
| *tetR*-$P_{tet}$ amplification | F_tetR_(ser) | R_tetR |
| $P_{tet}$ at the $tRNA^{Ser}$ locus | UF_tRNAser | R_tetR |
| *hlpA-mKate2* amplification (for $P_{32}$ fusion) | F_hlpA_(P32) | R_mKate2 |
| *hlpA-mNeonGreen* amplification (for $P_{32}$ fusion) | F_hlpA_(P32) | R_neonGreen |
| *hlpA-mScarlet-I* amplification (for $P_{32}$ fusion) | F_hlpA_(P32) | R_Scarlet |
| *hlpA-mTurquoise2* amplification (for $P_{32}$ fusion) | F_hlpA_(P32) | R_mTurquoise |
| *hlpA-msfYFP* amplification (for $P_{32}$ fusion) | F_hlpA_(P32) | R_YFP |
| *hlpA-mNeonGreen* at the $tRNA^{Ser}$ locus (for $P_{tet}$ fusion) | F_hlpA_(Ptet) | DR_tRNAser2 |
| *hlpA-mScarlet-I* at the $tRNA^{Ser}$ locus (for $P_{tet}$ fusion) | F_hlpA_(Ptet) | DR_tRNAser2 |
| *hlpA-mTurquoise2* at the $tRNA^{Ser}$ locus (for $P_{tet}$ fusion) | F_hlpA_(Ptet) | DR_tRNAser2 |
| *mNeonGreen* at the $tRNA^{Ser}$ locus | F_linker-fp | DR_tRNAser2 |
| *mScarlet-I* at the $tRNA^{Ser}$ locus | F_linker-fp | DR_tRNAser2 |
| *mTurquoise2* at the $tRNA^{Ser}$ locus | F_linker-fp | DR_tRNAser2 |
| *ftsZ* amplification (for $P_{tet}$ fusion) | F_ftsZ_(Ptet) | R_ftsZ_linkerfp |
| *divIVA* amplification (for $P_{32}$ fusion) | F_divIVA_(P32) | R_divIVA_linkerfp |
| *divIVA* at the $tRNA^{Ser}$ locus (for $P_{xyl1/2}$ fusion) | F_DivIVA_(Pxyl) | DR_tRNAser2 |
| *tetR*-$P_{tet}$-*ftsZ-mNeonGreen* amplification | F_tetR_(thr) | R_nGreen_(cat) |
| *xylR*-$P_{xyl1}$-*divIVA-mScarlet-I* amplification | F_xylR_GOR | R_fp_(ery) |
